# *Wolbachia* load variation in *Drosophila* is more likely caused by drift than by host genetic factors

**DOI:** 10.1101/2020.11.29.402545

**Authors:** Alexis Bénard, Hélène Henri, Camille Noûs, Fabrice Vavre, Natacha Kremer

## Abstract

Symbiosis is a continuum of long-term interactions ranging from mutualism to parasitism, according to the balance between costs and benefits for the protagonists. The density of endosymbionts is, in both cases, a key factor that determines both the transmission of symbionts and the host extended phenotype, and is thus tightly regulated within hosts. However, the evolutionary and molecular mechanisms underlying bacterial density regulation are currently poorly understood. In this context, the symbiosis between the fruit fly and its intracellular bacteria *Wolbachia* (*w*MelPop strain) is particularly interesting to study. Although vertically transmitted, the symbiont is pathogenic, and a positive correlation between virulence and *w*MelPop density is observed. In addition, the number of repeats of a bacterial genomic region -Octomom-is positively correlated with *Wolbachia* density, underlying a potential genetic mechanism that controls bacterial density. Interestingly, the number of repeats varies between host individuals, but most likely also within them. Such genetic heterogeneity within the host could promote conflicts between bacteria themselves and with the host, notably by increasing within-host competition between symbiont genotypes through a process analogous to the tragedy of the commons. To characterize the determinisms at play in the regulation of bacterial density, we first introgressed wMelPop in different genetic backgrounds of *D. melanogaster*, and found different density levels and Octomom copy numbers in each host lineage. To determine whether such variations reflect a host genetic determinism on density regulation through Octomom copy number selection, we replicated the introgressions and performed reciprocal crosses on the two Drosophila populations with the most extreme density levels. In both experiments, we detected an absence of directionality in the patterns of infection, associated with a strong instability of these patterns across generations. Given that bacterial density was highly correlated with Octomom copy numbers in all experiments, these results rather suggest a strong influence of drift and a random increase in the frequency of certain bacterial variants. We then discuss how drift, both on the symbiont population during transmission and on the host population, could limit the efficiency of selection in such a symbiotic system, and the consequences of drift on the regulation of density and composition of bacterial populations.

## Introduction

A majority of organisms live in symbiosis, a close relationship between two organisms belonging to different species that ranges along the continuum between parasitism and mutualism (De Bary, 1879; Tipton, Darcy and Hynson, 2019). In the case of microorganisms, the regulation of the symbiont population within the host, and particularly their abundance within host tissues, are important characteristics that shape the tight relationship between partners and influence the position of the symbiosis along the mutualism-parasitism continuum (Tiivel, 1991; Douglas, 1994). Research on disease evolution has further shown that the evolution of virulence is balanced by the transmission of symbionts to new hosts, and that both virulence and transmission rely on the regulation of the symbiotic density (Anderson and May, 1982). On the one side, an increased virulence can benefit symbionts by increasing their instantaneous transmission, as they exploit more host resources and thus increase their replication within the host. On the other side, the more abundant the symbionts are in host tissues, the more they cost to the host, which shortens the host life span and thereby the window of transmission of the symbiont. As a result, the virulence/transmission trade-off leads to a reproduction rate optimum that optimizes symbiont transmission over the entire life of the host. More specifically in vertically transmitted symbioses, the optimum symbiotic density optimizes both the production of offspring and their colonization by symbionts.

Symbiont density is thus under strong regulation (O’Neill, Hoffmann and Werren, 1997; Alizon *et al*., 2009), and many factors can contribute to its control (López-Madrigal and Duarte, 2019). In insects for instance, host factors can play a major role in regulating the symbiont population (Poinsot *et al*., 1998; Douglas, 2014) through the activation of immune pathways, such as DUOX or Toll (Douglas, Bouvaine and Russell, 2011; You, Lee and Lee, 2014). Symbionts can also be involved in their own regulation according to particular genetic factors (Ijichi *et al*., 2002; Chrostek *et al*., 2013). This is for example the case in symbioses between wasps and vertically transmitted bacteria, where densities of *Wolbachia* are strain-specific in co-infection (Mouton *et al*., 2003, 2004). Still, some mechanisms involved in bacterial regulation are poorly understood in insects. For instance, the target of bacterial regulation remains to be clarified: does the host control the overall symbiont population by decreasing symbiont abundance regardless the symbiont genetic specificity or does it target specific variants? Also, control mechanisms that are independent of classical immune pathways are worth exploring. For instance, are cases where hosts sanction symbiont through differential allocation of metabolites frequent and widespread in symbiotic associations (Douglas, 2008)?

There is much evidence to suggest that selection should lead to symbiotic population control systems (Douglas, 2014), but two evolutionary mechanisms could limit the effectiveness of selection on density regulation and should also be taken into consideration: conflicts between different levels of selection and drift. In terms of selection levels, between-host selection predicts that any excessive replication would be detrimental to host fitness, thus selecting for symbiotic variants that are the least harmful while being well transmitted (Szathmáry and Smith, 1995). On the contrary, the competition that occurs within the host tissues should favor symbiont variants that are the most efficient to rapidly colonize the host, thus those that have the most proliferative abilities regardless of the cost paid by the host (Alizon, de Roode and Michalakis, 2013). This raises the question of whether within- and between-host selection create an evolutionary conflict regarding the control of symbiont density, by favoring symbiont strains with opposite replication profiles (O’Neill, Hoffmann and Werren, 1997; Monnin *et al*., 2020). Finally, the importance of drift in vertically transmitted symbioses could be more considered. Indeed, bottlenecks during transmission reduce the genetic diversity in the following host generation and may limit the effectiveness of selection upon symbiotic population regulation (Mathé-Hubert *et al*., 2019). Such molecular and evolutionary mechanisms remain poorly studied, especially in vertically transmitted symbioses, although they can play an important role in the epidemiological and evolutionary dynamics of symbiotic interactions. A first limitation is conceptual, as populations of vertically transmitted endosymbionts tend to be considered with little or no heterogeneity, thus limiting the potential for within-host selection. However, while recurrent bottlenecks during transmission tend to reduce diversity, heterogeneity can still be observed in certain systems (Banks and Birky, 1985; Birky, Fuerst and Maruyama, 1989; Abbot and Moran, 2002; Asnicar *et al*., 2017). A second -more practical-limitation is that if heterogeneity does exist in symbiont populations, it is difficult to trace it experimentally, because of the absence of genetic markers.

A good study model to address questions related to density control is the maternally transmitted bacterium *Wolbachia* in association with *Drosophila melanogaster* hosts. In particular, the virulent *w*MelPop strain (Min and Benzer, 1997), which can exhibit heterogeneous density levels between individuals, has differential virulence profiles. Virulence is notably correlated to a tandem amplification of the genomic region “Octomom” (Chrostek *et al*., 2013). Indeed, flies harboring *Wolbachia* with more copies of Octomom exhibit high density levels in their tissues and a reduced lifespan, while those harboring *Wolbachia* with fewer copies exhibit low density levels and survive longer (Chrostek and Teixeira, 2015). This model system is therefore advantageous because hosts and symbionts can exhibit genetic variability, and because the number of Octomom copies can be used as a marker to track the evolution of the symbiotic population. Moreover, previous studies showed that within-host selection can occur in the *w*MelPop in *D. melanogaster* (Chrostek & Teixeira, 2018; Monnin et al. 2020).

In this study, we take advantage of this *Drosophila*-*w*MelPop symbiosis to shed light on the evolutionary determinisms that act on the regulation of vertically transmitted symbionts in insects. We investigate whether the host genetic background can directly influence the density of the symbionts, or whether the symbionts self-regulate their density *via* Octomom. Using different host genetic backgrounds and a combination of introgressions and crossing experiments, we analyze the respective role of host and symbiont backgrounds, but also drift, in the evolution of density and genetic composition of the symbiotic population.

## Methods

### Model system

*Drosophila melanogaster* flies were trapped in different locations (Arabia, Bolivia, China (Canton), Republic of the Congo - RC (Brazzaville), USA (Seattle) (Vieira et al., 1999) and France (Sainte-Foy-lès-Lyon)). These populations have been maintained in the laboratory by regular sib mating for at least 10 years and are considered as genetically homogeneous. In the following experiments, we used these 6 inbred lines (*Wolbachia*-free) plus the *w*1118 line, infected either by the *Wolbachia* strain *w*MelPop (provided by Scott O’Neill (Monash University, Australia)) or by the strain *w*MelCS (provided by J. Martinez/F. Jiggins, Cambridge University, UK).

### Rearing and collection

Flies were maintained under 12-hour day/night cycles at constant temperature and hygrometry (25°C and 60% relative humidity) and reared on rich medium (for 1 L of medium: 73.3 g of Gaude flour, 76.7 g of inactive brewer’s yeast, 8.89 g of agar-agar powder, 4 g of Tegosept - Nipagine, 0.4 L of distilled water and 55.5 mL of 95% ethanol). With the exception of introgression experiments (see below), each new generation was established by tube transfer of approximately 80 randomly selected 4/5-day old individuals, to ensure full fertility of the flies. To control larval competition prior to sampling for infection patterns, we pooled about 80 flies in egg-laying cages and transferred 100 eggs laid by 4/5-day old females onto a rich medium pellet (1 mL) placed in a tube of agarose medium. After hatching, flies were transferred onto an agarose medium supplemented with sugar (10 %) and were collected after 7 days to be frozen and stored at -20°C.

### *Wolbachia* introgression within various host genetic backgrounds

The symbiotic introgression method allows to transmit symbionts from a donor line to a recipient line while conserving most of the genetic background of the recipient line. As *Wolbachia* is a maternally transmitted bacterial symbiont, this method consists here in making a first cross between *Wolbachia*-infected females (here, n = 20) from the donor line with *Wolbachia*-uninfected males (here, n = 10) from the recipient line. Then, the F1 progeny of this previous cross carries the *Wolbachia* symbionts from the donor line and shares half of its genetic background between the donor and the recipient lines. Two additional backcrosses between females (n = 20) from the F1 (and then F2) progeny and males (n = 10) from the recipient lines are necessary to restore at the F3 generation 87.5 % of the genetic background of the recipient line (Figure 1).

**Figure 1:**
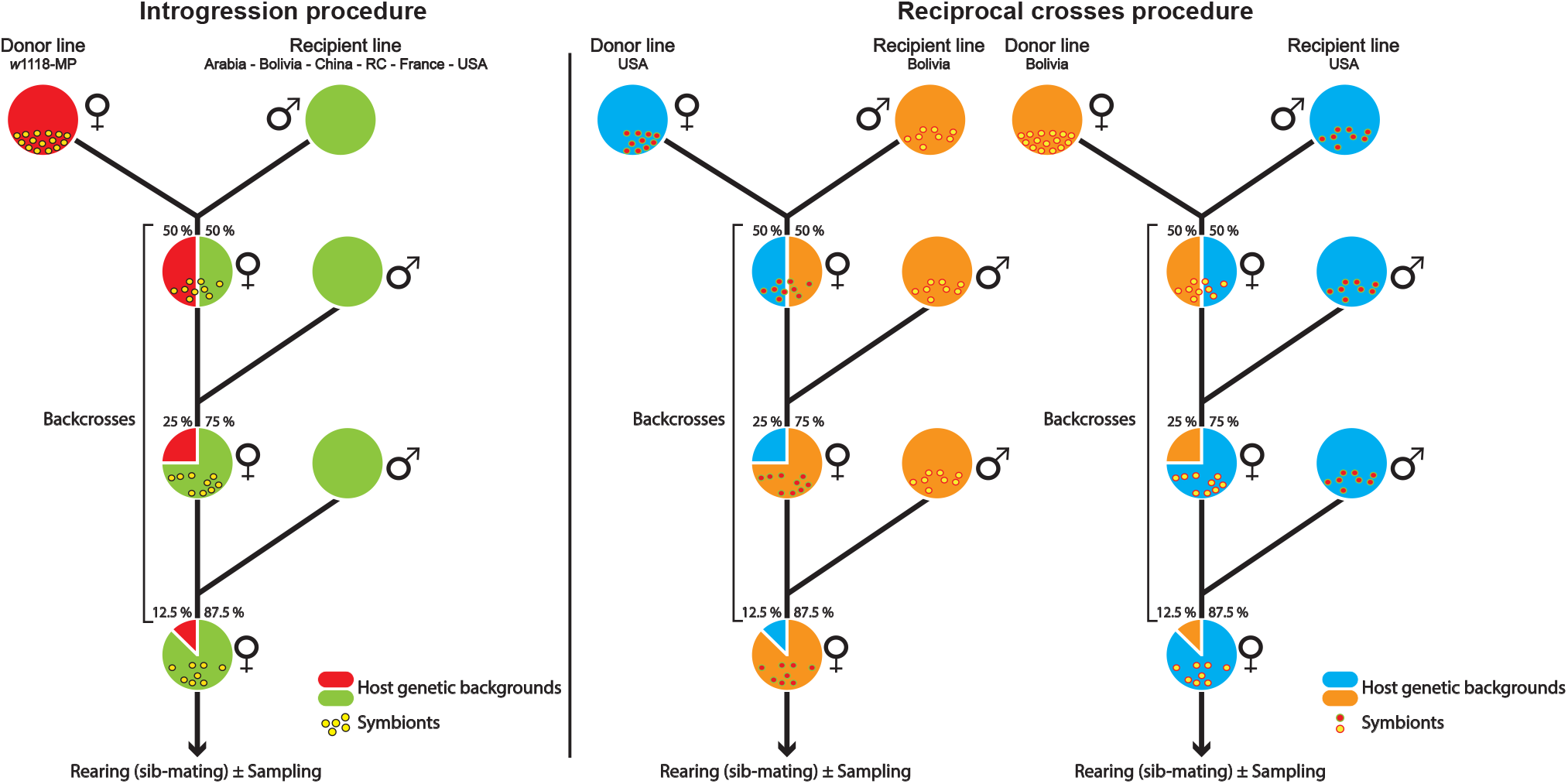
Introgression and reciprocal crosses procedures. Transmission of symbionts from females of the donor line to a recipient line. Serial backcrosses were performed to restore the recipient host genetic background by mating daughters from the previous cross with males from the recipient line. This method was applied to infect the 6 natural *Drosophila melanogaster* populations by *w*Melpop (experiment #1, left panel), to perform new introgressions from Bolivia or USA on 3 replicates (experiment #2, left panel) and to conduct reciprocal crosses (experiment #3, right panel).

We first applied this method to infect the 6 natural *Drosophila melanogaster* population lines by the *w*MelPop strain (experiment #1, MP1 lines). For this purpose, we used an iso-female *w*1118 line infected by *w*MelPop (IsoA3) as the donor line and the other populations as recipient lines (1 introgression / line). After two generations of regular sib-mating, flies were placed in egg-laying cages for sampling (see ‘rearing and collection’ protocol above), and infection patterns (*i*.*e*., *w*MelPop relative density and the average Octomom copy number per bacteria in flies) were checked by qPCR in these lines.

As the introgression of *w*MelPop in different recipient lines (experiment #1) resulted in different infection patterns (*i*.*e*., density and number of Octomom copies), we tested 8 generations later (experiment #2) the replicability of the infection pattern after a new introgression procedure. For this purpose, we selected two recipient lines (USA and Bolivia) that exhibited extreme infection patterns after introgression (*i*.*e*., USA-MP1 exhibited a high *w*MelPop density whereas Bolivia-MP1 exhibited a low *w*MelPop density, see results), and performed anew 3 independent symbiotic introgressions, using the same iso-female line (IsoA3, 12 generations after the first introgression procedure) as the donor line and these two populations (USA and Bolivia) as recipient lines. After 3 generations of backcrosses, Bolivia-MP2 and USA-MP2 flies were maintained under regular sib-mating (except the generation preceding each sampling, for which the larval density was controlled as described above).

In parallel to experiment #2, we independently performed reciprocal crosses between the Bolivia-MP1 and USA-MP1 lines (*i*.*e*., lines infected by *w*MelPop during the first introgression experiment) to test the respective influence of host and symbiotic genetic backgrounds on the *w*MelPop proliferation within flies. For this purpose, we reciprocally backcrossed Bolivia-MP1 and USA-MP1 individuals for 3 generations (experiment #3; 3 independent replicates), 8 generations after experiment #1. After 3 backcrosses, flies were maintained under regular sib-mating (except the generation preceding each sampling, in which the larval density was controlled).

### Quantification of *w*MelPop density and Octomom copy number

*Wolbachia* density and Octomom copy number were measured on 7-day old females (n = 10 flies / line (experiment #1) and n = 5 flies / line / timepoint (experiments #2 and #3)), whose DNA was extracted using the EZ-10 96-well Plate Animal Genomic DNA® kit (Bio Basic). In brief, flies were individually crushed in 400 µL of lysis buffer by a sterile 5-mm stainless bead shacked by a TissueLyser® (Qiagen) for 30 s at 25 Hz. DNA was extracted following the instructions from the manufacturer, eluted in 100 µL of elution buffer and stored at - 20°C.

Relative *Wolbachia* density and Octomom copy number were quantified from the same DNA extract by quantitative real-time PCR using SYBR® green and following the MIQE guideline applied to DNA samples (Bustin *et al*., 2009). To quantify the average amount of *w*MelPop per fly, we used primers targeting a monocopy reference gene in the host (*RP49* in *Drosophila melanogaster*) and primers targeting a monocopy gene outside of the Octomom region in *Wolbachia* (*WD0505* in *w*MelPop). Then, we normalized the number of copies of *WD0505* by the number of copies of the reference gene *RP49* to estimate the relative density of *w*MelPop per fly (Monnin *et al*., 2020). To quantify the average Octomom copy number of the *w*MelPop population within a fly, we used primers targeting the same gene located outside the Octomom copy number in the *w*MelPop genome (*WD0505*) and primers targeting a gene inside the Octomom region (*WD0513*). Then, we normalized the number of copies of *WD0513* by the number of copies of *WD0505* to estimate the mean Octomom copy number of the *w*MelPop population per fly (Chrostek *et al*., 2013). The sequences of the primers used (synthesis by Eurogentec®) are available in the Table s1.

The PCR amplifications were performed on a CFX96® instrument (Bio-Rad), independently for each target gene. Four µL of a diluted DNA sample (1/25), 0.5 µL of each forward and reverse primer (10 µM) and 5 µL of SsoADV Universal SYBR® Green Supermix (Bio-Rad) were used, for a total volume of 10 µL. The reaction conditions for amplification were 95 °C for 3 min of preincubation, followed by 40 cycles of {95 °C for 10 s for denaturation, 60 °C for 10 s for hybridization and 68 °C for 15 s for elongation}. The mean primer efficiencies were calculated using 6 points (in duplicate) from a 10-fold dilution series (103 to 108 copies) of previously purified PCR products (Table s1). The cycle quantification (Cq) values were estimated by the regression method, and the mean Cq value between technical duplicates was used for the determination of individual DNA quantities (deviation between duplicates below 0.5 cycles).

### Statistical analyses

We used the R software (version 4.0.3) for all analyses (R Core Team, 2020). Density and Octomom copy number ratios were estimated and normalized from the Cq values using the EasyqpcR package (Le Pape, 2012), based on the qBase algorithms published by Hellemans et al. (2007), taking into account the efficiency of primers. We first used a control sample from an aliquoted DNA extract (w1118 line infected by the *Wolbachia w*MelCS strain) as a calibrator, to estimate the inter-plate variability. We took this variability into account to normalize data between plates using the EasyqpcR package and determined the quantity of *WD0505* relative to *RP49* and of *WD0513* relative to *WD0505*. In addition, as the *w*MelCS genome contains only one copy of Octomom, we confirmed that the Octomom copy number measured was close to one and set its values to exactly 1. We used this transformation of the calibrator value as a standardization for all the samples.

The relative density data were analyzed using general linear models. Normality and homoscedasticity were checked graphically. The data on Octomom copy number were analyzed with general linear models with a gamma distribution, as the distribution of this factor did not fit to a normal distribution. We confirmed graphically that the gamma distribution used in the model fitted to the Octomom copy number data with the package fitdistrplus (Delignette-Muller and Dutang, 2015). The significance of the factors in these models were checked graphically with confidence intervals and considering p-values.

In the first experiment, we focused on the overall effect of the host genetic background on the relative density and Octomom copy number. The host genetic background of the lineages was thus set as the explanatory variable. We used the method of contrasts with *p*-values adjusted by Tukey method to obtain the pairwise differences between the lineages for both the relative density and the Octomom copy number. To determine the potential role of Octomom in the control of the bacterial population, we estimated the correlation between the relative density and the Octomom copy number (log-transformed data) using a linear model with the average Octomom copy number set as an explanatory variable.

In the second and third experiments, we focused on the differences between replicates. Then, the replicate label was set as the explanatory factor. The statistical analyses were performed independently for the Bolivia and USA host genetic backgrounds. We also used the method of contrasts with p-values adjusted by Tukey method to obtain the pairwise differences between the replicates for both the relative density and the Octomom copy number. We finally estimated the correlation between the relative density and the Octomom copy number (log-transformed data) using a linear model with the average Octomom copy number set as an explanatory variable. We performed these correlation analyses: 1) for each genetic background, separately for each timepoint, and 2) for each replicate line, with all timepoint grouped.

## Results

To characterize the determinisms at play in the regulation of bacterial density, we first investigated if host genetic background can have an active influence on density levels. As the Octomom region is also involved in density regulation, we additionally tested the influence of its amplification on density levels and the potential interaction between host and bacterial genotypes on density levels.

### *Wolbachia* introgression within various *Drosophila melanogaster* lines is associated with contrasted infection patterns

During a preliminary experiment, we checked the infection status of six *D. melanogaster* populations with contrasted genotypes, introgressed with the heterogeneous strain *w*MelPop originating from the same isoA3 line. When we quantified the relative *Wolbachia* density (Figure 2A) and the average copy number of the genomic region Octomom (Figure 2B) after the introgression protocol, we found contrasted infection patterns in the different *D. melanogaster* lines tested.

**Figure 2:**
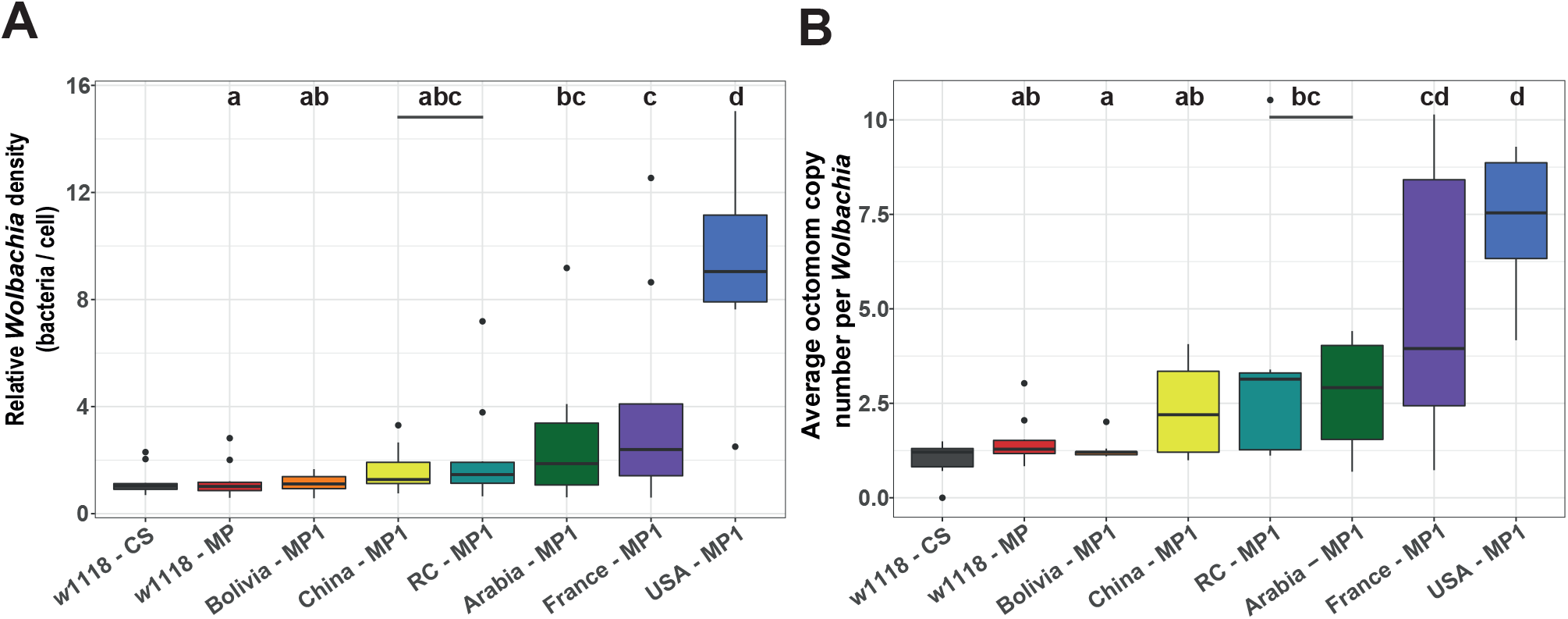
Infection patterns after introgression of *Wolbachia* in different host genetic backgrounds. **2A:** relative *Wolbachia* density per cell, **2B:** average Octomom copy number per *Wolbachia*. Each color represents a host genetic background (n = 10 flies / background). Box plots indicate ‘minimum’, 1^st^ quartile, median, 3^rd^ quartile, and ‘maximum’ ± outliers (dots). Different letters above boxplots indicate a significant difference between lines after pairwise comparisons (Table s3). The *w*1118-CS line is an experimental control infected by *w*MelCS and is not integrated in the statistical analyses. The *w*1118-MP line, infected by *w*MelPop, is the line initially used as ‘donor’ for the introgression procedure. All the other lines were infected by *w*MelPop by introgression (MP1).

Both *Wolbachia* density and composition (*i*.*e*., measured as the mean number of Octomom copies per *Wolbachia*) varied significantly among introgressed lines (Linear regression model; *w*1118 – MP; Population effect on relative density: *P* = 2.33 × 10^−09^ ; Population effect on Octomom copy number: *P* = 2.19 × 10^−08^ ; see statistical details in Table s2). Introgressed lines differed from each other (see pairwise comparisons in Table s3), with a maximum difference in bacterial density and mean Octomom copy number per *Wolbachia* of respectively 8.3 and 5.8-fold between Bolivia and USA lines (Table s2).

We observed a positive relationship between the relative density per line and the associated mean Octomom copy number per *Wolbachia* (*Intercept* = -0.40, *SE(intercept)* = 0.25, *slope* = 1.07, *SE(slope)* = 0.22, *r*^*2*^ = 0.83, Linear regression model on the median of each host genetic background : *P* = 0.005; Figure 3). This strong correlation suggests that variation in the number of Octomom copies is a genetic mechanism involved in the control of bacterial density and confirms previous results highlighted by Chrostek *et al*. (2013). The number of Octomom repeats could thus provide a way to monitor the evolution of bacterial populations across generations and to better characterize selective pressures associated with the control of bacterial populations.

**Figure 3:**
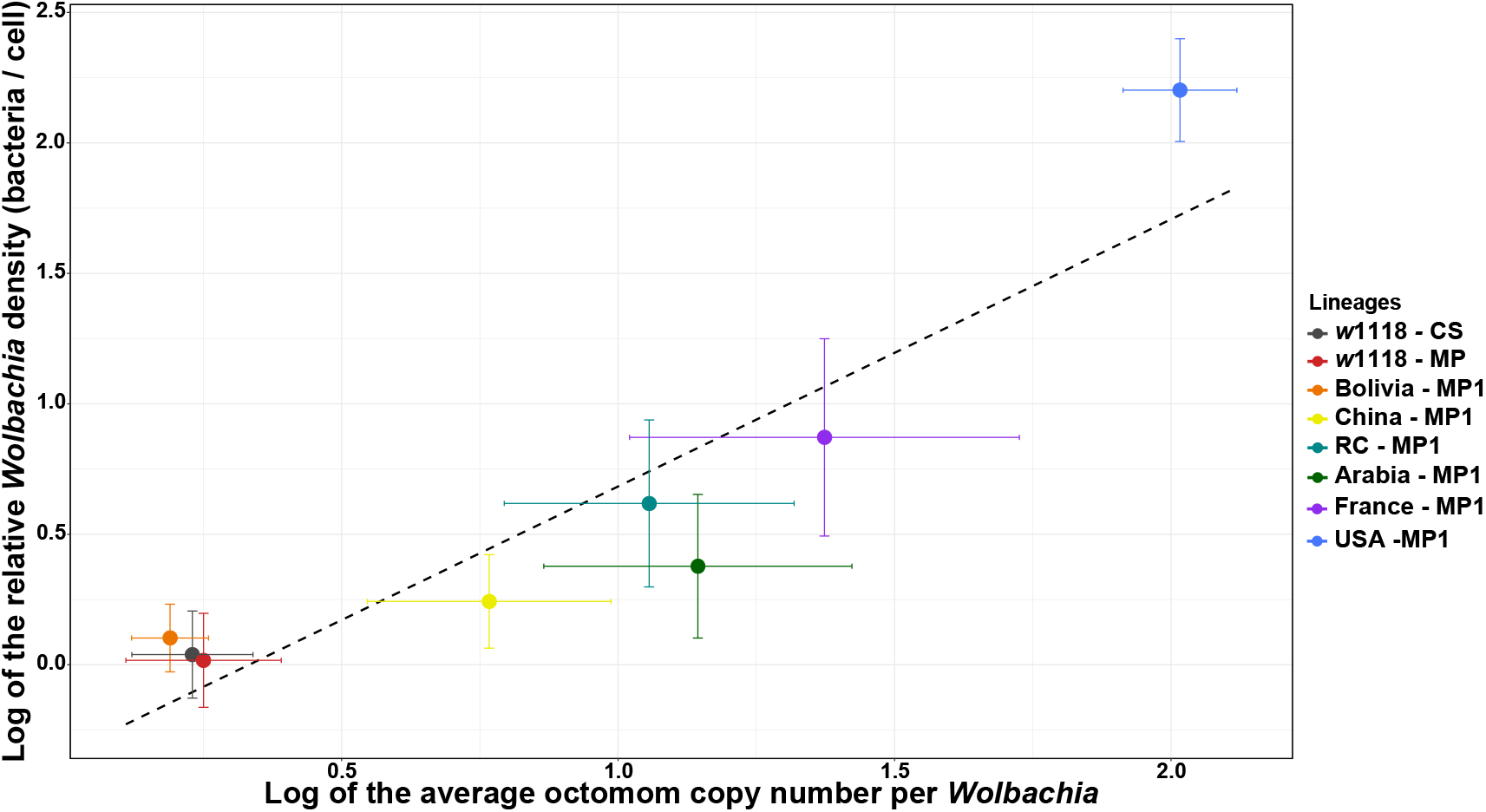
Relationship between the relative *w*MelPop density (log) and the average Octomom copy number per *Wolbachia* cell (log). Median ± SE. Each color represents a host genetic background (n = 10 flies / background), and the dashed line represents the linear regression.

To summarize, we observed in this preliminary experiment a large variation of *Wolbachia* densities between the six lines of *Drosophila melanogaster* introgressed with the same *w*MelPop line – densities that were strongly correlated to the average Octomom copy number per *Wolbachia*. These results are consistent with the selection of specific variants by different host genetic backgrounds. However, other factors, like genetic drift by founder effect during the vertical transmission of symbionts from the donor line and / or from one host generation to another, could explain this pattern. To disentangle these hypotheses, we thus performed two sets of experiments using the two lines that exhibited the most extreme patterns of infection in the preliminary experiment (*i*.*e*., Bolivia and USA).

### The infection pattern can change rapidly over generations, regardless of the host genetic background

In the first set of experiments, we performed a similar introgression of the *w*MelPop *Wolbachia* strain in the Bolivia and USA genetic backgrounds, but established three independent replicate lines for each host background (Bolivia-MP2 and USA-MP2 lines). While the three replicates should exhibit the same response under host control, variation among the three replicates is expected under the drift hypothesis. We additionally evaluated the stability of the infection pattern over generations, by tracking the relative *Wolbachia* density (Figure 4A) and the average Octomom copy number per *Wolbachia* (Figure 4B) immediately after the first introgression event, after 13 generations, and after 25 generations (see Tables s4 & s5 for details).

**Figure 4:**
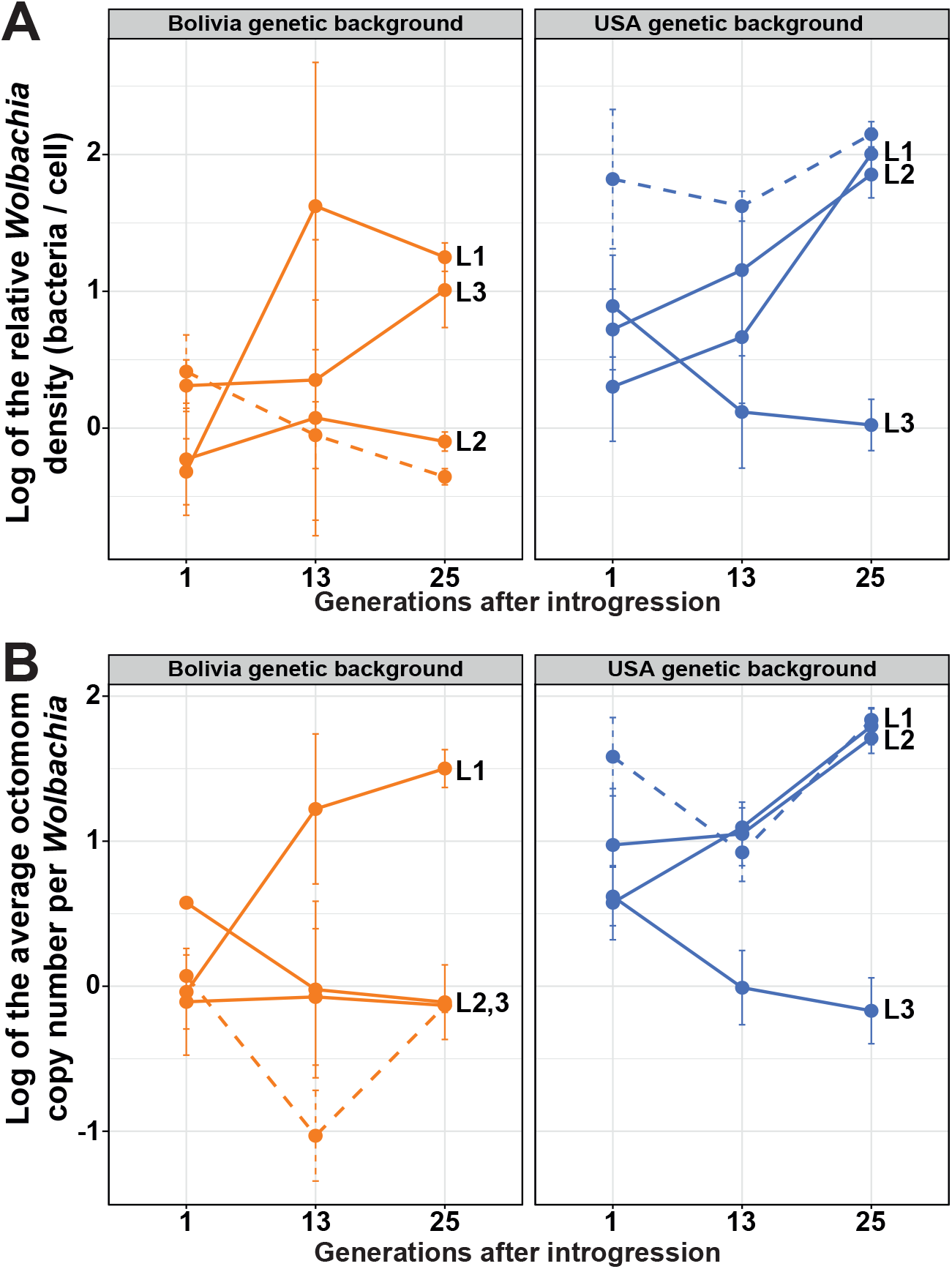
Replicability of the infection patterns after a new introgression procedure, and their evolution over generations. **4A:** relative *Wolbachia* density (log; median ± SE), **4B:** average Octomom copy number per *Wolbachia* (log; median ± SE), n = 5 / line / timepoint. Each color represents a host genetic background. Plain lines represent the replicate lineages from the new introgression procedure (MP2), L1, L2 and L3 indicating the replicates. Dashed lines represent the lineages from the initial introgression procedure (MP1), which were set as references in the statistical analyses.

Just after the introgression event (t = 1), the relative *Wolbachia* density and the average Octomom copy number per *Wolbachia* did not differ significantly between the Bolivia-MP2 replicate lines and the Bolivia-MP1 line from the first experiment used here as the reference (Linear regression model; experiment group effect on relative density: *P* > 0.1; experiment group effect on Octomom copy number: *P* > 0.1; see statistical details in Table s4). The relative density and Octomom copy number in *Wolbachia* from the Bolivia-MP2 replicate lines did not differ significantly between replicates (see pairwise comparisons in Table s5), which does not contradict a host determinism for density regulation trough Octomom copy number selection in this genetic background. However, at the same timepoint (t = 1), the relative density and the average Octomom copy number in *Wolbachia* from the USA-MP2 replicate lines tend to differ from the USA-MP1 line from the first experiment used here as the reference (Linear regression model; experiment group effect on relative density: *P* = 0.08; experiment group effect on Octomom copy number: *P* = 0.025; see statistical details in Table s4). The relative density and the average Octomom copy number in *Wolbachia* from the USA-MP2 replicates lines did not differ between replicates (see pairwise comparisons in Table s5) and the number of octomom copies per *Wolbachia* did not show significant difference with the donor line (w1118-MP) (Linear regression model; maternal transmission effect on Octomom copy number: *P* = 0.35; Table s4). These results are more consistent with a maternal transmission effect from the donor line to the recipient ones, with infection patterns mirroring the bacterial composition of the donor line.

We then examined the stability of the infection pattern over generations, for each replicate line (Figure 4). After 25 generations post introgression, the relative density and Octomom copy number per *Wolbachia* in Bolivia-MP2 and USA-MP2 replicate lines differed significantly from the quantities measured immediately after introgression (Linear regression model; generational effect on relative density: *P*_*Bolivia*_ = 6.22× 10^−06^, *P*_*USA*_ = 6.23× 10^−05^ ; generational effect on Octomom copy number: *P*_*Bolivia*_ = 7.25× 10^−04^, *P*_*USA*_ = 0.005; see statistical details in Table s4). Moreover, the infection patterns between the Bolivia-MP2 or between the USA-MP2 replicate lines significantly differed (see pairwise comparisons in Table s5).

Finally, we noticed that the correlation between the bacterial density and the average number of Octomom copies per *Wolbachia* remained high over generations (R^2^ > 71%, Figure s2) and within lines, when variation in *Wolbachia* density was observed (Figure s3 and associated statistics).

All together, these results show an absence of host control on the density and composition of the bacterial population, and an influence of the number of Octomom copies on the control of *Wolbachia* densities. The variations observed between replicate lines and over time may thus reflect drift.

### The bacterial composition initially transmitted strongly influences the patterns of infection observed over generations

In the second set of experiments, we performed reciprocal crosses to test the respective influence of the host genetic background, the bacterial population, the maternal effect of transmission, and drift on the *w*MelPop proliferation within flies (experiment #3). In order to jumble the host-*Wolbachia* associations, we made reciprocal crosses in 3 independent replicates using the Bolivia-MP1 and USA-MP1 lines from the first experiment, 8 generations after the latter. Then, to evaluate the stability of the infection pattern over generations, we measured the relative density of *w*MelPop (Figures 5A & 5B) and the average Octomom copy number per *Wolbachia* per fly (Figures 5C & 5D) one generation after the final homogenizing cross, 13 generations and 25 generations post introgression (see Tables s6 & s7 for details).

**Figure 5:**
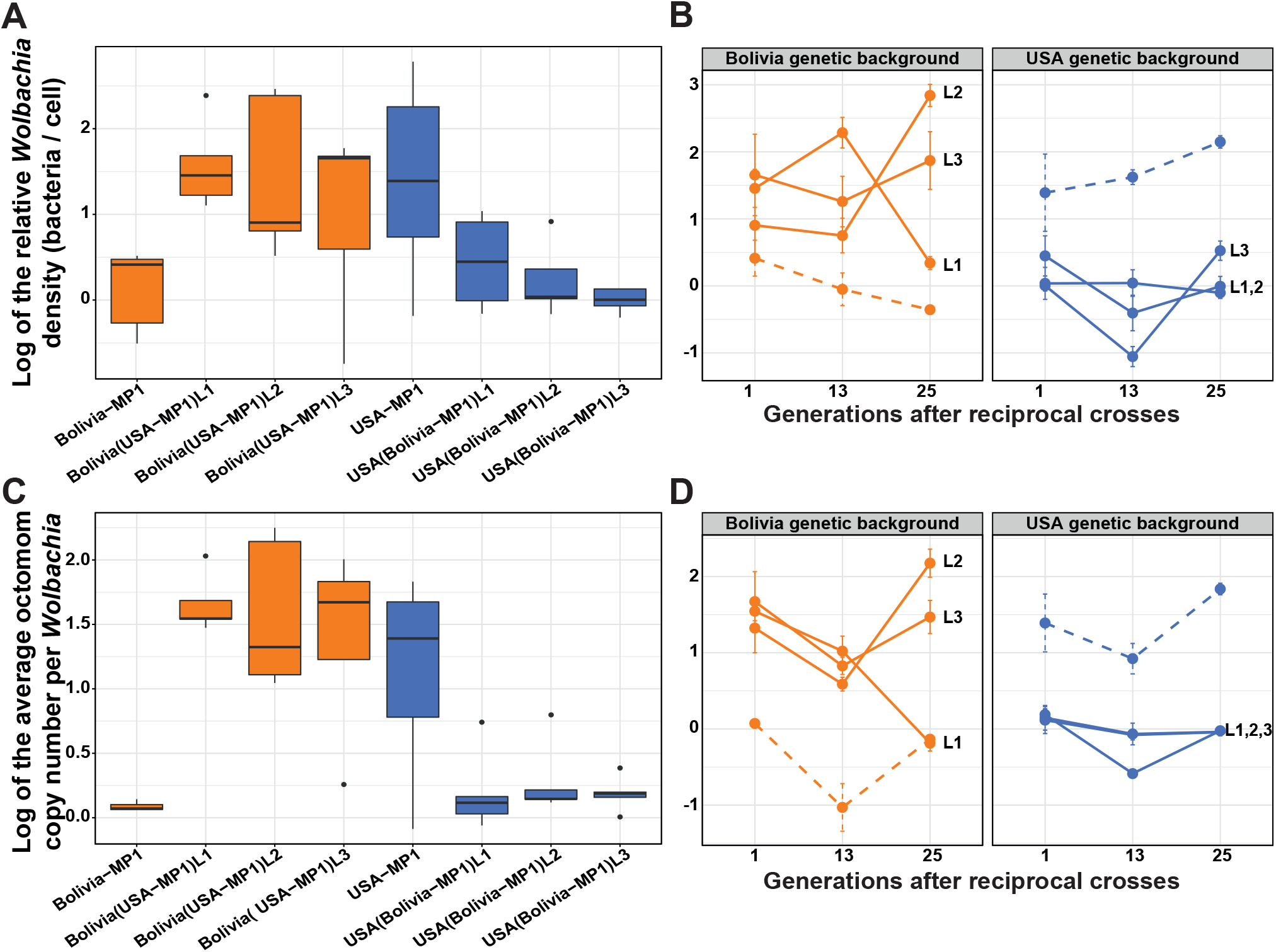
Evolution of infection patterns after reciprocal crosses. **5A:** Relative density (log) one generation post introgression (box plot with ‘minimum’, 1^st^ quartile, median, 3^rd^ quartile, and ‘maximum’ ± outliers (dots)). **5B:** Evolution of the relative density (log) over generations (median ± SE). **5C**: Average Octomom copy number per *Wolbachia* (log) one generation post introgression. **5D:** Evolution of the average Octomom copy number per *Wolbachia* (log) over generations (median ± SE). N = 5 flies / line / timepoint. Each color represents a host genetic background and the information in brackets represents the bacterial genetic background. Plain lines represent the replicate lineages from the reciprocal crosses, L1, L2 and L3 indicating the replicates. Dashed lines represent the lineages from the initial introgression procedure (MP1), which were used as references in the statistical analyses.

Just after the introgression event (t = 1), the relative density and the average Octomom copy number per *Wolbachia* differed significantly between the Bolivia(USA-MP1) replicate lines and the Bolivia-MP1 line (Linear regression model, experiment group effect on relative density: *P* = 0.047; experiment group effect on Octomom copy number: *P* = 4.88 × 10^−09^, see statistical details in Table s6), but not from the USA-MP1 line (Linear regression model, experiment group effect on relative density: *P* = 0.798; experiment group effect on Octomom copy number: *P* = 0.586, see statistical details in Table s6). In addition, the infection patterns of the Bolivia (USA-MP1) replicate lines did not differ significantly between them (see pairwise comparisons in Table s7). Similarly at the same timepoint, the relative density and the average Octomom copy number per *Wolbachia* from the USA(Bolivia-MP1) replicate lines differed significantly from those from the USA-MP1 line (Linear regression model, experiment group effect on relative density *P* = 0.018; experiment group effect on Octomom copy number *P* = 6.31 × 10^−08^, see statistical details in Table s6), but not from the Bolivia-MP1 line (Linear regression model, experiment group effect on relative density: *P* = 0.416; experiment group effect on Octomom copy number: *P* = 0.559, see statistical details in Table s6). In addition, the infection patterns of the USA(Bolivia-MP1) replicate lines did not differ significantly between them (see pairwise comparisons in Table s7). These results confirm an absence of control from the host on the establishment of the infection pattern and rather suggest a homogeneous symbiont transmission from the donor line.

Next, we examined whether infection patterns were stable within each of the replicate lines over generations by measuring the relative density and average Octomom copy number per *Wolbachia* 13 and 25 generations after the last backcross. After 25 generations post introgression, the relative density in Bolivia(USA-MP1) replicate lines differed significantly from their reference at t = 1 (Linear regression model, generational effect on the relative density: *P* = 0.015; generational effect on the Octomom copy number: *P* = 7.79 × 10^−05^, see statistical details in Table s6). On the contrary, the USA(Bolivia-MP1) replicate lines did not differ from their reference at t = 1 (Linear regression model, generational effect on the relative density: *P* = 0.052; generational effect on the Octomom copy number: *P* = 0.059, see statistical details in Table s6). Moreover, the infection patterns of the Bolivia(USA-MP1) replicate lines differed significantly from each other, as did the infection patterns of the USA(Bolivia-MP1) replicate lines (see pairwise comparisons in Table s7). All together, these results confirm an absence of host control on the density and composition of the bacterial population. Moreover, the variability observed between the replicates indicates a random transmission of the symbiont over generations.

Finally, we confirmed that the correlation between the bacterial density and the average number of Octomom copies per *Wolbachia* remained high over generations (R^2^ > 83%, Figure s4) and within the lines, when variation in *Wolbachia* density was observed (Figure s5 and associated statistics).

In conclusion, we failed to reveal any influence of the host genotype on the control of the *w*MelPop proliferation through the selection of bacteria containing high or low Octomom copy number. Instead, we found a strong maternal effect of transmission and an instability of the infection patterns over generations. These experiments lead us to consider that drift could be an important evolutionary force responsible for the diversification of infection patterns observed.

## Discussion

In this study we sought to identify the determinisms involved in the regulation of endosymbiotic populations and used the *Drosophila melanogaster* – *w*MelPop symbiotic system to track the influence of host and symbiont genotypes on the density regulation, as well as the evolutionary forces at play. Indeed, this symbiotic model is particularly relevant because it exhibits genetic variability among the population of vertically transmitted symbionts, whose evolution can be tracked by a genomic amplification (Octomom). While we first found large differences when comparing infection patterns (*i*.*e*., bacterial density and average Octomom copy number per *Wolbachia*) in different host genetic backgrounds, such host control on bacterial proliferation/selection was not confirmed after new experiments of introgression on more replicate lines and crossing experiments between lines exhibiting the most extreme infection patterns. Instead, we showed that the infection patterns were initially set up by the bacterial genotype and became very unstable over generations. These results suggest that, in this symbiotic system and under these experimental conditions, drift strongly influences the evolution of the symbiont density -and thus its stability over generations-, contrarily to what is generally described in the literature of insect endosymbioses (*e*.*g*., Mouton et al., 2003; Hosokawa *et al*., 2006).

Numerous examples in insects support an active regulation of symbiotic populations by the host, with stable density over generations when the environment remains constant (Ikeda, Ishikawa and Sasaki, 2003; Mouton *et al*., 2004, 2007; Funkhouser-Jones *et al*., 2018). The orchestrated modulation of the symbiont proliferation rate throughout insect development also suggests a fine-tuned host control of the bacterial density (Rio *et al*., 2006; Login *et al*., 2011; Vigneron *et al*., 2014). On the opposite, bacterial factors alone can also explain variation in bacterial densities within some hosts. For example, different strains of *Wolbachia* are known to exhibit different, but stable, density levels in the same host lines based on their genetic particularities (Mouton *et al*., 2003; Chrostek *et al*., 2013). Proliferation of symbionts within the host is under strong selection as it is a key factor influencing the trade-off between symbiont transmission (*i*.*e*., the higher the symbiont density, the higher the probability of transmission) and virulence (*i*.*e*., the higher the symbiont density, the higher the cost on host survival and fecundity) (Anderson and May, 1982; Ewald, 1983). This transmission/virulence trade-off often leads to an optimal density, which can be controlled by host or bacterial determinants. In insect hosts, the main molecular mechanisms that determine the abundance and composition of symbionts are associated with immune response (Lemaitre and Hoffmann, 2007; Zug and Hammerstein, 2015) or resource allocation (Kiers *et al*., 2003). Microbial communities can in turn select resistance mechanisms against host effectors or trigger antagonist regulators of the host immunity (Vallet-Gely *et al*., 2008; Lindsey, 2020).

In the *D. melanogaster-w*MelPop system, however, we observed a strong instability of infection patterns. Because the introgression was limited to 87.5%, a few alleles from the donor background – different in each replicate line – could marginally influence the *Wolbachia* load. However, incomplete introgression should not affect the results of the experiment where lines with the two most extreme phenotypes were crossed. Alternatively, the high instability could be due either to an instability of the optimum, or to a large influence of drift that limits the ability of the system to reach the optimum. Our results, and notably the variations observed between replicate lines in controlled rearing conditions, rather suggest a strong influence of drift on the regulation of bacterial density. Bacterial factors, such as the number of Octomom copies, could fluctuate through time and be at the origin of variation in density levels. However, the genomic region ‘Octomom’ has recently been questioned regarding its involvement in the establishment of density levels. Initially, it has been shown that proliferation rate and virulence of *w*MelPop are correlated with genomic amplification of the Octomom region (Chrostek *et al*., 2013; Chrostek and Teixeira, 2015, 2018), but this relationship has been challenged by Rohrscheib and her collaborators (Rohrscheib *et al*., 2016, 2017), who support that the virulence of *w*MelPop rather depends on an increase in the extrinsic rearing temperature. However, to exclude any influence of the Octomom copy number on *Wolbachia* growth and pathogenicity, these variables should be tested independently of the temperature (Chrostek and Teixeira, 2017). At constant temperature, our results show a clear link between *Wolbachia* density in adults and the average number of Octomom copies per *Wolbachia*, and are in accordance with the current literature (Chrostek *et al*., 2013; Chrostek and Teixeira, 2015, 2018; Monnin *et al*., 2020). While we cannot exclude that another gene or set of *Wolbachia* genes different from Octomom could determine symbiont density, the strong correlation between the density observed and the number of Octomom repeats in all the experiments (generally >80% when the density was variable) suggests that Octomom is the main determinant of bacterial density in this biological system and in our controlled conditions. Consequently, this genomic amplification can be used as a marker of bacterial diversity and evolution of our experimental system.

We can thus wonder why we observed such variability and temporal instability within lineages, and why drift overcame this potential bacterial regulation through Octomom? Indeed, this pattern contrasts with what is observed in already well-established symbioses, where one symbiont genotype is fixed (Werren, Baldo and Clark, 2008). In our experiments, we were able to show very similar levels of infection between mothers and daughters just after introgression or reciprocal crosses procedures, suggesting a maternal effect. However, instability detected across generations suggests that this maternal effect is probably non-genetic: when a large number of bacteria is quantified in the mother’s tissues, a large number of bacteria is transmitted to the oocytes and maintained in the adult stage (Veneti *et al*., 2004; Hosokawa, Kikuchi and Fukatsu, 2007; Parkinson, Gobin and Hughes, 2016). However, there may still be random variability between mothers regarding the amount of bacteria transmitted to their eggs, and between these eggs (Mira and Moran, 2002). Bottlenecks during transmission can thus eventually lead to a gradual shift of the ‘initial’ density over time. Bottlenecks can also influence density levels through random differential transmission of bacterial variants from one generation to the next (Funk, Wernegreen and Moran, 2001; Kaltenpoth *et al*., 2010), especially if these variants exhibit different reproductive rates (as it is the case with variants carrying different numbers of Octomom copies (Duarte *et al*., 2021)). To summarize, if not counteracted by host or symbiont density control, drift is expected to induce instability over generations by a combination of quantitative (*i*.*e*., transmission of a non-equivalent number of bacteria to the eggs) and qualitative/genetic (*i*.*e*., random transmission of different variants) bottlenecks. Hence, the high variability and the temporal instability depicted in our study could reflect the random transmission of different *w*MelPop quantities and variants during transmission bottlenecks. In addition, drift at the transmission level alone cannot explain variation of the average density at the population level but should be associated to other factors. Drift at the host population level could play a role by fixing hosts with symbiotic populations with a different average of Octomom copy number. Alternatively, selection, or mutation bias could also play a role. For instance, the mutation rate of a repeated sequence in microsatellites can strongly depend on the number of motifs present in the sequence (Whittaker *et al*., 2003).

Assuming the same rules on the number of Octomom copies present in the population (*i*.*e*., a higher propensity for duplication when the number of copies is high), the outcomes of drift could be more unpredictable for individuals harboring the highest average number of Octomom copies. Indeed, in the presence of moderate bottlenecks and a mutation rate increasing with the number of copies, a strong variability of infection patterns is expected over generations. However, we did not observe such higher temporal instability when the values of density and Octomom copy numbers were high in the donor lines. These results suggest that the influence of the mutation rate was negligible compared to the transmission bottleneck. Under conditions of genetic instability linked to transmission bottlenecks, between-host selection should not be efficient, and would explain why the vertically transmitted *w*MelPop strain exhibits a strong virulence, whereas the overall alignment of interests between the host and vertically transmitted symbionts generally leads to the selection of low virulent symbionts that maximize host survival and indirectly their own transmission (Anderson and May, 1982; O’Neill, Hoffmann and Werren, 1997).

Different environmental conditions can also modify the optimum density of the symbiont. Optima can be different for the host and the symbionts, and lead to antagonistic interactions between symbiotic partners and to variations in bacterial density (Parker *et al*., 2021). In the *D. melanogaster* / *w*MelPop system, the maintenance of the virulence phenotype has thus frequently been associated with the fact that virulence is only expressed in conditions rarely observed in nature, so that the between-host selection against highly prolific variants (such as those with high Octomom copy numbers) is weak at 25°C. A recent study however shows that strains with 8-9 Octomom copies are pathogenic from 18°C to 29°C (Duarte *et al*., 2021). In addition, another selective force, the within-host selection, could explain the virulence of *w*MelPop in certain environmental conditions. Indeed, when fly populations are reared at 28°C, Monnin *et al*. (2020) showed that the population evolved toward a higher virulence, which may be due to the stronger effect of within-host selection compared to between-host selection. Thus, when selective pressures are strong, within and between-host selection could modulate symbiont virulence in the *Drosophila*-*w*MelPop association, whereas drift might not allow any co-evolution between partners and co-adaptation to environmental changes when selective pressures are limited.

To conclude, we showed that the host did not control for bacterial density and composition in the symbiosis between *D. melanogaster* and *w*MelPop, and that the infection patterns were very instable across generations, suggesting a strong influence of drift that could limit the effects of within- and between-host selections. As the transmission of symbionts in vertically transmitted symbiosis is subject to potential bottlenecks both in terms of quantity and genetic diversity (Mira and Moran, 2002; Galbreath *et al*., 2009; Kaltenpoth *et al*., 2010), it seems necessary to further characterize the intensity of bottlenecks in this symbiotic system, in order to better evaluate the impact of drift on the evolution of bacterial populations in vertically transmitted symbioses and its impact on host phenotypes.

## Data accessibility

Raw data are available online: https://doi.org/10.5281/zenodo.4607210

## Supplementary material

Additional figures are available online: https://www.biorxiv.org/content/10.1101/2020.11.29.402545v4.supplementary-material

Scripts are available online: https://doi.org/10.5281/zenodo.4607223

## Acknowledgements

We thank Nicole Lara and Claire Traversaz-Erchoff for providing the Drosophila media. We also thank the DTAMB platform for providing us technical resources for molecular biology experiments. This work was supported by the Agence Nationale de la Recherche ‘RESIST’ program (ANR-16-CE02-0013-01) and performed within the framework of the LABEX ECOFECT (ANR-11-LABX-0048) and the SBP Micro-Be-Have of Université de Lyon, within the programme ‘Investissements d’Avenir’ (ANR-11-IDEX-0007; ANR-16-IDEX-0005). Version 4 of this preprint has been peer-reviewed and recommended by Peer Community In Evolutionary Biology (https://doi.org/10.24072/pci.evolbiol.100126).

## Conflict of interest disclosure

The authors of this preprint declare that they have no financial conflict of interest with the content of this article. NK & FV are recommenders for PCI Evolutionary Biology.

## Notes

https://doi.org/10.5281/zenodo.4607210

https://doi.org/10.5281/zenodo.4607223

### Competing Interest Statement

The authors have declared no competing interest.

### Summary of Updates

Version 4 of this preprint has been peer-reviewed and recommended by Peer Community In Evolutionary Biology (https://doi.org/10.24072/pci.evolbiol.100126)

## References

Abbot, P. and Moran, N. A. (2002) ‘Extremely low levels of genetic polymorphism in endosymbionts (Buchnera) of aphids (Pemphigus)’, Molecular Ecology, 11(12), pp. 2649–2660. doi: 10.1046/j.1365-294X.2002.01646.x.

Alizon, S. et al. (2009) ‘Virulence evolution and the trade-off hypothesis: History, current state of affairs and the future’, Journal of Evolutionary Biology, 22(2), pp. 245–259. doi: 10.1111/j.1420-9101.2008.01658.x.

Alizon, S., de Roode, J. C. and Michalakis, Y. (2013) ‘Multiple infections and the evolution of virulence’, Ecology Letters, 16(4), pp. 556–567. doi: 10.1111/ele.12076.

Anderson, R. M. and May, R. M. (1982) ‘Coevolution of hosts and parasites’, Parasitology. Cambridge University Press, 85(2), pp. 411–426. doi: 10.1017/S0031182000055360.

Asnicar, F. et al. (2017) ‘Studying Vertical Microbiome Transmission from Mothers to Infants by Strain-Level Metagenomic Profiling’, mSystems, 2(1), pp. e00164–16. doi: 10.1128/mSystems.00164-16.

Banks, J. A. and Birky, C. W. (1985) ‘Chloroplast DNA diversity is low in a wild plant, Lupinus texensis.’, Proceedings of the National Academy of Sciences of the United States of America, 82(20), pp. 6950–6954. doi: 10.1073/pnas.82.20.6950.

Birky, C. W., Fuerst, P. and Maruyama, T. (1989) ‘Organelle gene diversity under migration, mutation, and drift: Equilibrium expectations, approach to equilibrium, effects of heteroplasmic cells, and comparison to nuclear genes’, Genetics, 121(3), pp. 613–627. doi: 10.1111/j.1365-2028.1973.tb00101.x.

Bustin, S. A. et al. (2009) ‘The MIQE guidelines: Minimum information for publication of quantitative real-time PCR experiments’, Clinical Chemistry, 55(4), pp. 611–622. doi: 10.1373/clinchem.2008.112797.

Chrostek, E. et al. (2013) ‘Wolbachia Variants Induce Differential Protection to Viruses in Drosophila melanogaster: A Phenotypic and Phylogenomic Analysis’, PLoS Genetics, 9(12). doi: 10.1371/journal.pgen.1003896.

Chrostek, E. and Teixeira, L. (2015) ‘Mutualism Breakdown by Amplification of Wolbachia Genes’, PLoS Biology, 13(2), pp. 1–23. doi: 10.1371/journal.pbio.1002065.

Chrostek, E. and Teixeira, L. (2017) ‘Comment on Rohrscheib et al. 2016 ‘Intensity of mutualism breakdown is determined by temperature not amplification of Wolbachia genes’, PLoS Pathogens, 13(9), pp. 2–7. doi: 10.1371/journal.ppat.1006521.

Chrostek, E. and Teixeira, L. (2018) ‘Within host selection for faster replicating bacterial symbionts’, PLoS ONE, 13(1), pp. 1–8. doi: 10.1371/journal.pone.0191530.

De Bary, A. (1879) ‘Die Erscheinung der Symbiose’. Vortrag auf der Versammlung der Naturforshung und Ärtze zu Cassel. Verlag von Karl J. Trübner Strassburg, 121, pp. 1–30.

Delignette-Muller, M. L. and Dutang, C. (2015) ‘{fitdistrplus}: An {R} Package for Fitting Distributions’, Journal of Statistical Software, 64(4), pp. 1–34. Available at: http://www.jstatsoft.org/v64/i04/.

Douglas, A. E. (1994) Symbiotic interactions. Oxon, GB: Oxford University Press, 1994.

Douglas, A. E. (2008) ‘Conflict, cheats and the persistence of symbioses’, New Phytologist, 177(4), pp. 849–858. doi: 10.1111/j.1469-8137.2007.02326.x.

Douglas, A. E. (2014) ‘The molecular basis of bacterial-insect symbiosis’, Journal of Molecular Biology. Elsevier B.V., 426(23), pp. 3830–3837. doi: 10.1016/j.jmb.2014.04.005.

Douglas, A. E., Bouvaine, S. and Russell, R. R. (2011) ‘How the insect immune system interacts with an obligate symbiotic bacterium’, Proceedings of the Royal Society B: Biological Sciences, 278(1704), pp. 333–338. doi: 10.1098/rspb.2010.1563.

Duarte, E. H. et al. (2021) ‘Forward genetics in Wolbachia: Regulation of Wolbachia proliferation by the amplification and deletion of an addictive genomic island’, bioRxiv. doi:10.1101/2020.09.08.288217

Ewald, P. W. (1983) ‘Host-Parasite relations, vectors, and the evolution od disease severity’, Annual Review of Ecology, Evolution, and Systematics, 14, pp. 465–485.

Funk, D. J., Wernegreen, J. J. and Moran, N. A. (2001) ‘Intraspecific variation in symbiont genomes: bottlenecks and the aphid-Buchnera association’, Genetics, 157(2), pp. 477–489.

Funkhouser-Jones et al. (2018) ‘The maternal effect gene Wds controls Wolbachia titer in Nasonia’, Current Biology, 28(11), pp.1692–1702.e6. doi:10.1016/j.cub.2018.04.010.

Galbreath, J. G. M. S. et al. (2009) ‘Reduction in post-invasion genetic diversity in Crangonyx pseudogracilis (Amphipoda: Crustacea): A genetic bottleneck or the work of hitchhiking vertically transmitted microparasites?’, Biological Invasions, 12(1), pp. 191–209. doi: 10.1007/s10530-009-9442-3.

Hellemans, J. et al. (2007) ‘qBase relative quantification framewok and software for management and automated analysis of real-time quantitative PCR data’, Genome Biology, 8:R19. doi:10.1186/gb-2007-8-2-r19.

Hosokawa, T. et al. (2006). ‘Strict Host-Symbiont cospeciation and reductive genome evolution in insect gut bacteria’, PLoS Biology, 4(10), pp. e337. doi:10.1371/journal.pbio.0040337.

Hosokawa, T., Kikuchi, Y. and Fukatsu, T. (2007) ‘How many symbionts are provided by mothers, acquired by offspring, and needed for successful vertical transmission in an obligate insect-bacterium mutualism?’, Molecular Ecology, 16(24), pp. 5316–5325. doi: 10.1111/j.1365-294X.2007.03592.x.

Ijichi, N. et al. (2002) ‘Internal Spatiotemporal Population Dynamics of Infection with Three Wolbachia Strains in the Adzuki Bean Beetle, Callosobruchus chinensis (Coleoptera: Bruchidae)’, American Society for Microbiology, 68(8), pp. 4074–4080. doi: 10.1128/AEM.68.8.4074.

Ikeda, T., Ishikawa, H. and Sasaki, T. (2003) ‘Regulation of Wolbachia Density in the Mediterranean Flour Moth, Ephestia kuehniella, and the Almond Moth, Cadra cautella’, Zoological Science, 20(2), pp. 153–157. doi: 10.2108/zsj.20.153.

Kaltenpoth, M. et al. (2010) ‘Life cycle and population dynamics of a protective insect symbiont reveal severe bottlenecks during vertical transmission’, Evolutionary Ecology, 24(2), pp. 463–477. doi: 10.1007/s10682-009-9319-z.

Kiers, E.T., Rousseau, R.A., West, S.A., Denison, R.F. (2003) ‘Host sanctions and the legume-rhizobium mutualism’, Nature, 425, pp. 78–81. doi:10.1038/nature01931.

Lemaitre, B. and Hoffmann, J. (2007) ‘The Host Defense of Drosophila melanogaster’, Annual Review of Immunology, 25, pp. 697–743. doi: 10.1146/annurev.immunol.25.022106.141615.

Le Pape, S. (2012) ‘EasyqpcR: EasyqpcR for easy analysis of real-time PCR data at IRTOMIT-INSERM U1082’. Available at: http://irtomit.labo.univ-poitiers.fr/.

Lindsey, A.R.I. (2020) ‘Sensing, signaling, and secretion: a review and analysis of systems for regulating host interaction in Wolbachia’, Genes 11(7):813. doi: 10.3390/genes11070813.

Login, F. H. et al. (2011) ‘Antimicrobial peptides keep insect endosymbionts under control’, Science, 334(6054), pp. 362–365. doi: 10.1126/science.1209728.

López-Madrigal, S., and Duarte, E. H. (2019) ‘Titer regulation in arthropod-Wolbachia symbioses’, FEMS Microbiology Letters, 366(23), fnz232. doi: 10.1093/femsle/fnz232

Mathé-Hubert, H. et al. (2019) ‘Nonrandom associations of maternally transmitted symbionts in insects: The roles of drift versus biased cotransmission and selection’, Molecular Ecology, 28(24), pp. 5330–5346. doi: 10.1111/mec.15206.

Min, K. T. and Benzer, S. (1997) ‘Wolbachia, normally a symbiont of Drosophila, can be virulent, causing degeneration and early death.’, Proceedings of the National Academy of Sciences of the United States of America, 94(20), pp. 10792–10796. doi: 10.1073/pnas.94.20.10792.

Mira, A. and Moran, N. A. (2002) ‘Estimating population size and transmission bottlenecks in maternally transmitted endosymbiotic bacteria’, Microbial Ecology, 44(2), pp. 137–143. doi: 10.1007/s00248-002-0012-9.

Monnin, D. et al. (2020) ‘Experimental evolution of virulence and associated traits in a Drosophila melanogaster – Wolbachia symbiosis’, bioRxiv, 2020.04.26.062265, ver. 4 peer-reviewed and recommended by PCI Evol Biol. doi: https://doi.org/10.1101/2020.04.26. 062265.

Mouton, L. et al. (2003) ‘Strain-specific regulation of intracellular Wolbachia density in multiply infected insects’, Molecular Ecology, 12(12), pp. 3459–3465. doi: 10.1046/j.1365-294X.2003.02015.x.

Mouton, L. et al. (2004) ‘Virulence, multiple infections and regulation of symbiotic population in the Wolbachia-Asobara tabida symbiosis’, Genetics, 168(1), pp. 181–189. doi: 10.1534/genetics.104.026716.

Mouton, L. et al. (2007) ‘Interaction between host genotype and environmental conditions affects bacterial density in Wolbachia symbiosis’, Biology Letters, 3(2), pp. 210–213. doi: 10.1098/rsbl.2006.0590.

O’Neill, S. L., Hoffmann, A. and Werren, J. (1997) Influential passengers: inherited microorganisms and arthropod reproduction. Oxford University Press.

Parker, B. J, et al. (2021) ‘Intraspecific variation in symbiont density in an insect-microbe symbiosis’, Molecular Ecology, doi: 10.1111/mec.15821.

Parkinson, J. F., Gobin, B. and Hughes, W. O. H. (2016) ‘Heritability of symbiont density reveals distinct regulatory mechanisms in a tripartite symbiosis’, Ecology and Evolution, 6(7), pp. 2053–2060. doi: 10.1002/ece3.2005.

Poinsot, D. et al. (1998) ‘Wolbachia Transfer from Drosophila melanogaster into D. simulans: Host Effect and Cytoplasmic Incompatibility Relationships’, Genetics, 150(1), pp. 227–237.

Rio, R. V. M. et al. (2006) ‘Dynamics of multiple symbiont density regulation during host development: Tsetse fly and its microbial flora’, Proceedings of the Royal Society B: Biological Sciences, 273(1588), pp. 805–814. doi: 10.1098/rspb.2005.3399.

Rohrscheib, C. E. et al. (2016) ‘Intensity of Mutualism Breakdown Is Determined by Temperature Not Amplification of Wolbachia Genes’, PLoS Pathogens, 12(9), pp. 1–15. doi: 10.1371/journal.ppat.1005888.

Rohrscheib, C. E. et al. (2017) ‘Response to: Comment on Rohrscheib et al. 2016 “Intensity of mutualism breakdown is determined by temperature not amplification of Wolbachia genes”’, PLoS Pathogens, 13(9), pp. 2016–2018. doi: 10.1371/journal.ppat.1006521.

Szathmáry, E. and Smith, J. M. (1995) ‘The major evolutionary transitions’, Nature, pp. 227–232. doi: 10.1038/374227a0.

Tiivel, T. (1991) ‘Cell Symbiosis, Adaptation, and Evolution: Insect-Bacteria Examples’, in: Symbiosis as a Source of Evolutionary Innovation: Speciation and Morphogenesis. MIT Press, p. 170.

Tipton, L., Darcy, J. L. and Hynson, N. A. (2019) ‘A developing symbiosis: Enabling cross-talk between ecologists and microbiome scientists’, Frontiers in Microbiology, 10(292). doi: 10.3389/fmicb.2019.00292.

Vallet-Gely, I., Lemaitre, B., Boccard, F. (2008) ‘Bacterial strategies to overcome insect defences’, Nature Reviews Immunology 6, pp.302–313. doi: doi:10.1038/nrmicro1870.

Veneti, Z. et al. (2004) ‘Heads or tails: Host-parasite interactions in the Drosophila-Wolbachia system’, Applied and Environmental Microbiology, 70(9), pp. 5366–5372. doi: 10.1128/AEM.70.9.5366-5372.2004.

Vieira, C. Lepetit, D., Dumont, S. and Biémont C. (1999) ‘Wake up of transposable elements following Drosophila simulans worldwide colonization’, Mol. Biol. Evol. 16(9):1251–55.

Vigneron, A. et al. (2014) ‘Insects Recycle Endosymbionts when the Benefit is Over’, Current Biology, 24, 2267–73. http://dx.doi.org/10.1016/j.cub.2014.07.065.

Werren, J.H., Baldo, L., and Clark, M.E. (2008) ‘Wolbachia: master manipulators of invertebrate biology’, Nature Reviews Microbiology, 6 pp.741–751. doi:10.1038/nrmicro1969.

Whittaker, J. C. et al. (2003) ‘Likelihood-based estimation of microsatellite mutation rates’, Genetics, 164(2), pp. 781–787.

You, H., Lee W. J., and Lee, W-J. (2014) ‘Homeostasis between gut-associated microorganisms and the immune system in Drosophila’, Current Opinion in Immunology, 30, pp.48–53. doi: 10.1016/j.coi.2014.06.006

Zug, R., and Hammerstein, P. (2015) ‘Wolbachia and the insect immune system: what reactive oxygen species can tell us about mechanisms of Wolbachia-host interactions’. Frontiers in Microbiology, 6:1201. doi: 0.3389/fmicb.2015.01201.

