## Supporting information for "*Wolbachia* load variation in *Drosophila* is more likely caused by drift than by host genetic factors"

Alexis Bénard [1,*], Hélène Henri [1], Camille Noûs [1,2], Fabrice Vavre [1], Natacha Kremer [1,*]

[1] Université de Lyon, Université Lyon 1, CNRS, VetAgroSup, Laboratoire de Biométrie et Biologie Evolutive, UMR 5558, F-69622 Villeurbanne, France.

[2] Laboratoire Cogitamus, France.

<u>**Table s1:**</u> **Characteristics of oligonucleotide primers** used for the quantification of *Wolbachia* relative density and average Octomom copy number by qPCR.

<u>**Table s2:**</u> **Statistics associated with linear regression models on data from experiment #1** (relative *w*MelPop density and average Octomom copy number).

<u>**Table s3:**</u> **Pairwise comparisons performed from linear regression outputs of the experiment #1** (relative *w*MelPop density and average Octomom copy number).

<u>**Table s4:**</u> **Statistics associated with linear regression models on data from experiment #2** (relative *w*MelPop density and average Octomom copy number).

<u>**Table s5:**</u> **Pairwise comparisons performed on the linear regression models from the second introgression experiment** (relative *w*MelPop density and average Octomom copy number).

<u>**Table s6:**</u> **Statistics associated with linear regression models on data from the reciprocal crosses experiment** (relative *w*MelPop density and average Octomom copy).

<u>**Table s7:**</u> **Pairwise comparisons performed on the linear regression models from the reciprocal crosses experiment** (relative *w*MelPop density and average Octomom copy).

<u>**Figure s1:**</u> **Relationship between the relative density and the average octomom copy number per *Wolbachia*** in replicate lines at each timepoint of experiment #1.

<u>**Figure s2:**</u> **Relationship between the relative density and the average octomom copy number per *Wolbachia*** in replicate lines at each timepoint of the new introgression experiment (exp. #2).

<u>**Figure s3:**</u> **Correlation between the relative density and the average octomom copy number per *Wolbachia*** within each replicate line of the new introgression experiment (exp. #2).

<u>**Figure s4:**</u> **Correlation between the relative density and the average octomom copy number per *Wolbachia*** in replicate lines at each timepoint of the 'reciprocal crosses' experiment (exp. #3).

<u>**Figure s5:**</u> **Correlation between the relative density and the average octomom copy number per *Wolbachia*** within each line of the 'reciprocal crosses' experiment (exp. #3).

**<u>Table s1:</u> Characteristics of oligonucleotide primers** used for the quantification of *Wolbachia* relative density and average Octomom copy number by qPCR. The amplification efficiency (mean ± SE) was determined by in-run standard curves, using 2 duplicates of a 10-fold dilution series of purified PCR products from $10^3$ to $10^8$ copies.

| Gene | Sequence | Efficiency | References |
|---|---|---|---|
| RP49dd-R<br>RP49dd-F | 5'-CGA-TCT-CGC-CGC-AGT-AAC-3'<br>5'-CTG-CCC-ACC-GGA-TTC-AAG-3' | 96.7 % ± 0.6 | Steckel and Boutros, 2005 |
| WD0505-qR<br>WD0505-qF | 5'-ACG-CGA-GCA-TCT-TCC-ATA-AG-3'<br>5'-TGT-TCC-TGG-TGG-ATC-ATC-TG-3' | 94.6 % ± 0.7 | Chrostek *et al.*, 2013 |
| WD0513-qR<br>WD0513-qF | 5'-AGC-ATG-TCC-TCT-GTG-CCA-TC-3'<br>5'-TTA-ACC-GGC-CAG-TCT-TAT-CG-3' | 93.3 % ± 0.9 | Chrostek *et al.*, 2013 |

Chrostek, E. *et al.* (2013) '*Wolbachia* Variants Induce Differential Protection to Viruses in *Drosophila melanogaster*: A Phenotypic and Phylogenomic Analysis', *PLoS Genetics*, 9(12). doi: 10.1371/journal.pgen.1003896.

Steckel, M. and Boutros, M. (2005) 'Rapid Development of Real-Time RT-PCR Assays Using Universal ProbeLibrary : Applications for Dissecting Signaling Pathways by RNA Interference', *Biochemica*, 3, pp. 17–19.

**<u>Table s2:</u> Statistics associated with linear regression models on data from experiment #1** (relative *w*MelPop density and average Octomom copy number per *Wolbachia*).

| Lines | Mean density | SEM density | Mean Octomom | SEM Octomom | *P*-value density | *P*-value Octomom |
|---|---|---|---|---|---|---|
| **w1118-MP** | 1.23 | 0.21 | 1.50 | 0.20 | Reference | Reference |
| **Arabia-MP1** | 2.69 | 0.81 | 2.78 | 0.45 | 0.047 | 0.047 |
| **Bolivia-MP1** | 1.14 | 0.11 | 1.27 | 0.08 | 0.955 | 0.633 |
| **RC-MP1** | 2.16 | 0.62 | 3.17 | 0.88 | 0.149 | 0.030 |
| **China-MP1** | 1.62 | 0.26 | 2.29 | 0.37 | 0.325 | 0.169 |
| **France-MP1** | 3.87 | 1.22 | 5.02 | 1.13 | 0.004 | < 0.001 |
| **USA-MP1** | 9.48 | 1.13 | 7.40 | 0.54 | < 0.001 | < 0.001 |
| **w1118-CS** | 1.20 | 0.17 | 1.03 | 0.14 | ND | ND |

**<u>Table s3:</u> Pairwise comparisons performed from linear regression outputs of the experiment #1.** Summary information of the pairwise comparisons with p-values adjusted using the Tukey's method. Analyses from bacterial density (log-transformed data) and Octomom copy numbers (log-transformed data) respectively below and above the black diagonal.

| | *w*1118-MP | Arabia-MP1 | Bolivia-MP1 | RC-MP1 | China-MP1 | USA-MP1 | France-MP1 |
|---|---|---|---|---|---|---|---|
| *w*1118-MP | | 0.147 | 0.993 | 0.340 | 0.589 | < 0.001 | < 0.001 |
| Arabia-MP1 | 0.041 | | 0.022 | 0.998 | 0.986 | 0.001 | 0.186 |
| Bolivia-MP1 | 1.000 | 0.372 | | 0.003 | 0.185 | < 0.001 | < 0.001 |
| RC-MP1 | 0.767 | 0.997 | 0.733 | | 0.834 | 0.009 | 0.491 |
| China-MP1 | 0.954 | 0.943 | 0.940 | 0.999 | | < 0.001 | 0.022 |
| USA-MP1 | < 0.001 | < 0.001 | < 0.001 | < 0.001 | < 0.001 | | 0.683 |
| France-MP1 | 0.005 | 0.961 | 0.050 | 0.728 | 0.427 | 0.001 | |

Octomom Copy Number

Bacterial density

**<u>Table s4:</u> Statistics associated with linear regression models on data from experiment #2** (relative *w*MelPop density and average Octomom copy number).

| Replicate lines | Mean density | SEM density | Mean Octomom | SEM Octomom | *P*-value density | *P*-value Octomom | Generation |
|---|---|---|---|---|---|---|---|
| ***w*1118-MP** | 1.01 | 0.14 | 1.72 | 0.23 | | | 1 |
| **Bolivia-MP1** | 1.51 | 0.29 | 1.07 | 0.02 | > 0.1 | > 0.1 | 1 |
| **Bolivia-MP2-L1** | 0.72 | 0.24 | 0.96 | 0.40 | | | |
| **Bolivia-MP2-L2** | 0.80 | 0.63 | 0.90 | 0.63 | | | |
| **Bolivia-MP2-L3** | 1.36 | 0.22 | 1.78 | 0.05 | | | |
| ***w*1118-MP *vs* Bolivia-MP2lines** | ND | ND | ND | ND | 0.699 | 0.718 | 1 |
| **Bolivia-MP1** | 0.95 | 0.36 | 0.36 | 0.22 | > 0.1 | 0.022 | 13 |
| **Bolivia-MP2-L1** | 5.07 | 8.20 | 3.40 | 1.49 | | | |
| **Bolivia-MP2-L2** | 1.07 | 7.32 | 0.93 | 1.23 | | | |
| **Bolivia-MP2-L3** | 1.42 | 8.40 | 0.98 | 1.84 | | | |
| **Bolivia-MP1** | 0.70 | 0.05 | 0.87 | 0.04 | < 0.001 | < 0.001 | 25 |
| **Bolivia-MP2-L1** | 3.49 | 0.42 | 4.49 | 0.66 | | | |
| **Bolivia-MP2-L2** | 0.91 | 0.07 | 0.88 | 0.03 | | | |
| **Bolivia-MP2-L3** | 2.75 | 0.52 | 0.90 | 0.23 | | | |
| **USA-MP1** | 6.17 | 3.36 | 4.86 | 0.98 | 0.081 | 0.025 | 1 |
| **USA-MP2-L1** | 1.35 | 1.02 | 1.78 | 0.44 | | | |
| **USA-MP2-L2** | 2.06 | 0.95 | 2.65 | 1.88 | | | |
| **USA-MP2-L3** | 2.44 | 1.22 | 1.86 | 0.36 | | | |
| ***w*1118-MP *vs* USA-MP2 lines** | ND | ND | ND | ND | 0.015 | 0.345 | 1 |
| **USA-MP1** | 5.06 | 0.47 | 2.51 | 0.47 | > 0.1 | 0.005 | 13 |
| **USA-MP2-L1** | 1.95 | 2.86 | 2.99 | 0.51 | | | |
| **USA-MP2-L2** | 3.18 | 2.73 | 2.86 | 0.65 | | | |
| **USA-MP2-L3** | 1.04 | 0.40 | 1.30 | 0.33 | | | |
| **USA-MP1** | 8.57 | 0.73 | 6.28 | 0.47 | < 0.001 | < 0.001 | 25 |
| **USA-MP2-L1** | 7.41 | 1.39 | 6.01 | 0.75 | | | |
| **USA-MP2-L2** | 6.39 | 0.85 | 5.53 | 0.49 | | | |
| **USA-MP2-L3** | 1.02 | 0.17 | 0.84 | 0.26 | | | |
| **Generation effect in Bolivia-MP2 lines** | ND | ND | ND | ND | < 0.001 | < 0.001 | 1 *vs.* 25 |
| **Generation effect in USA-MP2 lines** | ND | ND | ND | ND | < 0.001 | 0.005 | 1 *vs.* 25 |

**<u>Table s5:</u> Pairwise comparisons performed on the linear regression models from the second introgression experiment.**

Summary information of the pairwise comparison with adjusted P-values with Tukey's method (bacterial density and Octomom copy numbers respectively below and above the black diagonal).

| Generation 1 | Bolivia-MP2-L1 | Bolivia-MP2-L2 | Bolivia-MP2-L3 | USA-MP2-L1 | USA-MP2-L2 | USA-MP2-L3 |
|---|---|---|---|---|---|---|
| Bolivia-MP2-L1 | | 0.699 | 0.626 | ND | ND | ND |
| Bolivia-MP2-L2 | 0.513 | | 0.999 | ND | ND | ND |
| Bolivia-MP2-L3 | 0.548 | 0.999 | | ND | ND | ND |
| USA-MP2-L1 | ND | ND | ND | | 0.270 | 0.999 |
| USA-MP2-L2 | ND | ND | ND | 0.733 | | 0.304 |
| USA-MP2-L3 | ND | ND | ND | 0.727 | 1.000 | |

Octomom Copy Number (above diagonal); Bacterial density (below diagonal)

| Generation 13 | Bolivia-MP2-L1 | Bolivia-MP2-L2 | Bolivia-MP2-L3 | USA-MP2-L1 | USA-MP2-L2 | USA-MP2-L3 |
|---|---|---|---|---|---|---|
| Bolivia-MP2-L1 | | 0.831 | 0.999 | ND | ND | ND |
| Bolivia-MP2-L2 | 0.929 | | 0.799 | ND | ND | ND |
| Bolivia-MP2-L3 | 0.999 | 0.908 | | ND | ND | ND |
| USA-MP2-L1 | ND | ND | ND | | 0.921 | 0.013 |
| USA-MP2-L2 | ND | ND | ND | 0.996 | | 0.044 |
| USA-MP2-L3 | ND | ND | ND | 0.510 | 0.390 | |

Octomom Copy Number (above diagonal); Bacterial density (below diagonal)

| Generation 25 | Bolivia-MP2-L1 | Bolivia-MP2-L2 | Bolivia-MP2-L3 | USA-MP2-L1 | USA-MP2-L2 | USA-MP2-L3 |
|---|---|---|---|---|---|---|
| Bolivia-MP2-L1 | | < 0.001 | < 0.001 | ND | ND | ND |
| Bolivia-MP2-L2 | < 0.001 | | 0.943 | ND | ND | ND |
| Bolivia-MP2-L3 | 0.011 | 0.003 | | ND | ND | ND |
| USA-MP2-L1 | ND | ND | ND | | 0.806 | < 0.001 |
| USA-MP2-L2 | ND | ND | ND | 0.394 | | < 0.001 |
| USA-MP2-L3 | ND | ND | ND | < 0.001 | < 0.001 | |

Octomom Copy Number (above diagonal); Bacterial density (below diagonal)

**<u>Table s6:</u> Statistics associated with linear regression models on data from the reciprocal crosses experiment** (relative *w*MelPop density and average Octomom copy).

| Replicate lines | Mean density | SEM density | Mean Octomom | SEM Octomom | *P*-value density | *P*-value Octomom | Generation |
|---|---|---|---|---|---|---|---|
| **Bolivia-MP1** | 1.23 | 0.28 | 1.09 | 0.02 | 0.047 | < 0.001 | 1 |
| **Bolivia(USA-MP1)L1** | 5.39 | 1.79 | 5.34 | 0.74 | | | |
| **Bolivia(USA-MP1)L2** | 5.80 | 2.83 | 5.53 | 1.80 | | | |
| **Bolivia(USA-MP1)L3** | 3.75 | 1.37 | 4.74 | 1.36 | | | |
| **Bolivia-MP1** | 1.24 | 0.36 | 0.53 | 0.22 | < 0.001 | < 0.001 | 13 |
| **Bolivia(USA-MP1)L1** | 9.20 | 2.01 | 3.37 | 0.67 | | | |
| **Bolivia(USA-MP1)L2** | 2.26 | 0.64 | 1.91 | 0.17 | | | |
| **Bolivia(USA-MP1)L3** | 3.85 | 1.19 | 2.34 | 0.26 | | | |
| **Bolivia-MP1** | 0.73 | 0.04 | 0.87 | 0.04 | < 0.001 | < 0.001 | 25 |
| **Bolivia(USA-MP1)L1** | 1.51 | 0.14 | 0.83 | 0.08 | | | |
| **Bolivia(USA-MP1)L2** | 16.58 | 2.64 | 7.99 | 1.38 | | | |
| **Bolivia(USA-MP1)L3** | 10.99 | 4.54 | 4.47 | 0.93 | | | |
| **USA-MP1** | 6.46 | 3.09 | 3.79 | 1.08 | 0.018 | < 0.001 | 1 |
| **USA(Bolivia-MP1)L1** | 1.74 | 0.49 | 1.27 | 0.26 | | | |
| **USA(Bolivia-MP1)L2** | 1.37 | 0.37 | 1.38 | 0.26 | | | |
| **USA(Bolivia-MP1)L3** | 1.01 | 0.08 | 1.21 | 0.09 | | | |
| **USA-MP1** | 4.80 | 0.48 | 2.52 | 0.47 | < 0.001 | < 0.001 | 13 |
| **USA(Bolivia-MP1)L1** | 0.79 | 0.20 | 0.88 | 0.05 | | | |
| **USA(Bolivia-MP1)L2** | 1.01 | 0.17 | 1.07 | 0.15 | | | |
| **USA(Bolivia-MP1)L3** | 0.36 | 0.05 | 0.56 | 0.02 | | | |
| **USA-MP1** | 8.50 | 0.73 | 6.20 | 0.47 | < 0.001 | < 0.001 | 25 |
| **USA(Bolivia-MP1)L1** | 1.05 | 0.15 | 0.96 | 0.05 | | | |
| **USA(Bolivia-MP1)L2** | 0.94 | 0.08 | 0.99 | 0.04 | | | |
| **USA(Bolivia-MP1)L3** | 1.99 | 0.29 | 0.99 | 0.03 | | | |
| **Generation effect in Bolivia(USA-MP1) lines** | ND | ND | ND | ND | 0.015 | < 0.001 | 1 *vs.* 25 |
| **Generation effect in USA(Bolivia-MP1) lines** | ND | ND | ND | ND | 0.052 | 0.059 | 1 *vs.* 25 |
| **Bolivia(USA-MP1) lines *vs* USA-MP1** | ND | ND | ND | ND | 0.798 | 0.586 | 1 |
| **USA(Bolivia-MP1)lines *vs* Bolivia-MP1** | ND | ND | ND | ND | 0.416 | 0.559 | 1 |

**<u>Table s7:</u> Pairwise comparisons performed on the linear regression models from the 'reciprocal crosses' experiment.**

Summary information of the pairwise comparison with adjusted P-values with Tukey's method (bacterial density and Octomom copy numbers respectively below and above the black diagonal).

| Generation 1 | Bolivia(USA-MP1)L1 | Bolivia(USA-MP1)L2 | Bolivia(USA-MP1)L3 | USA(Bolivia-MP1)L1 | USA(Bolivia-MP1)L2 | USA(Bolivia-MP1)L3 |
|---|---|---|---|---|---|---|
| Bolivia(USA-MP1)L1 | | 0.999 | 0.966 | ND | ND | ND |
| Bolivia(USA-MP1)L2 | 0.989 | | 0.933 | ND | ND | ND |
| Bolivia(USA-MP1)L3 | 0.664 | 0.832 | | ND | ND | ND |
| USA(Bolivia-MP1)L1 | ND | ND | ND | | 0.988 | 0.998 |
| USA(Bolivia-MP1)L2 | ND | ND | ND | 0.962 | | 0.957 |
| USA(Bolivia-MP1)L3 | ND | ND | ND | 0.744 | 0.949 | |

Octomom Copy Number (above diagonal); Bacterial density (below diagonal)

| Generation 13 | Bolivia(USA-MP1)L1 | Bolivia(USA-MP1)L2 | Bolivia(USA-MP1)L3 | USA(Bolivia-MP1)L1 | USA(Bolivia-MP1)L2 | USA(Bolivia-MP1)L3 |
|---|---|---|---|---|---|---|
| Bolivia(USA-MP1)L1 | | 0.206 | 0.562 | ND | ND | ND |
| Bolivia(USA-MP1)L2 | 0.002 | | 0.882 | ND | ND | ND |
| Bolivia(USA-MP1)L3 | 0.037 | 0.482 | | ND | ND | ND |
| USA(Bolivia-MP1)L1 | ND | ND | ND | | 0.537 | 0.010 |
| USA(Bolivia-MP1)L2 | ND | ND | ND | 0.535 | | < 0.001 |
| USA(Bolivia-MP1)L3 | ND | ND | ND | 0.018 | 0.001 | |

Octomom Copy Number (above diagonal); Bacterial density (below diagonal)

| Generation 25 | Bolivia(USA-MP1)L1 | Bolivia(USA-MP1)L2 | Bolivia(USA-MP1)L3 | USA(Bolivia-MP1)L1 | USA(Bolivia-MP1)L2 | USA(Bolivia-MP1)L3 |
|---|---|---|---|---|---|---|
| Bolivia(USA-MP1)L1 | | < 0.001 | < 0.001 | ND | ND | ND |
| Bolivia(USA-MP1)L2 | < 0.001 | | 0.006 | ND | ND | ND |
| Bolivia(USA-MP1)L3 | < 0.001 | 0.145 | | ND | ND | ND |
| USA(Bolivia-MP1)L1 | ND | ND | ND | | 0.981 | 0.956 |
| USA(Bolivia-MP1)L2 | ND | ND | ND | 0.904 | | 0.999 |
| USA(Bolivia-MP1)L3 | ND | ND | ND | 0.001 | < 0.001 | |

Octomom Copy Number (above diagonal); Bacterial density (below diagonal)

<u>Figure s1:</u> **Relationship between the relative density and the average octomom copy number per *Wolbachia* in replicate lines at each timepoint of experiment #1.**

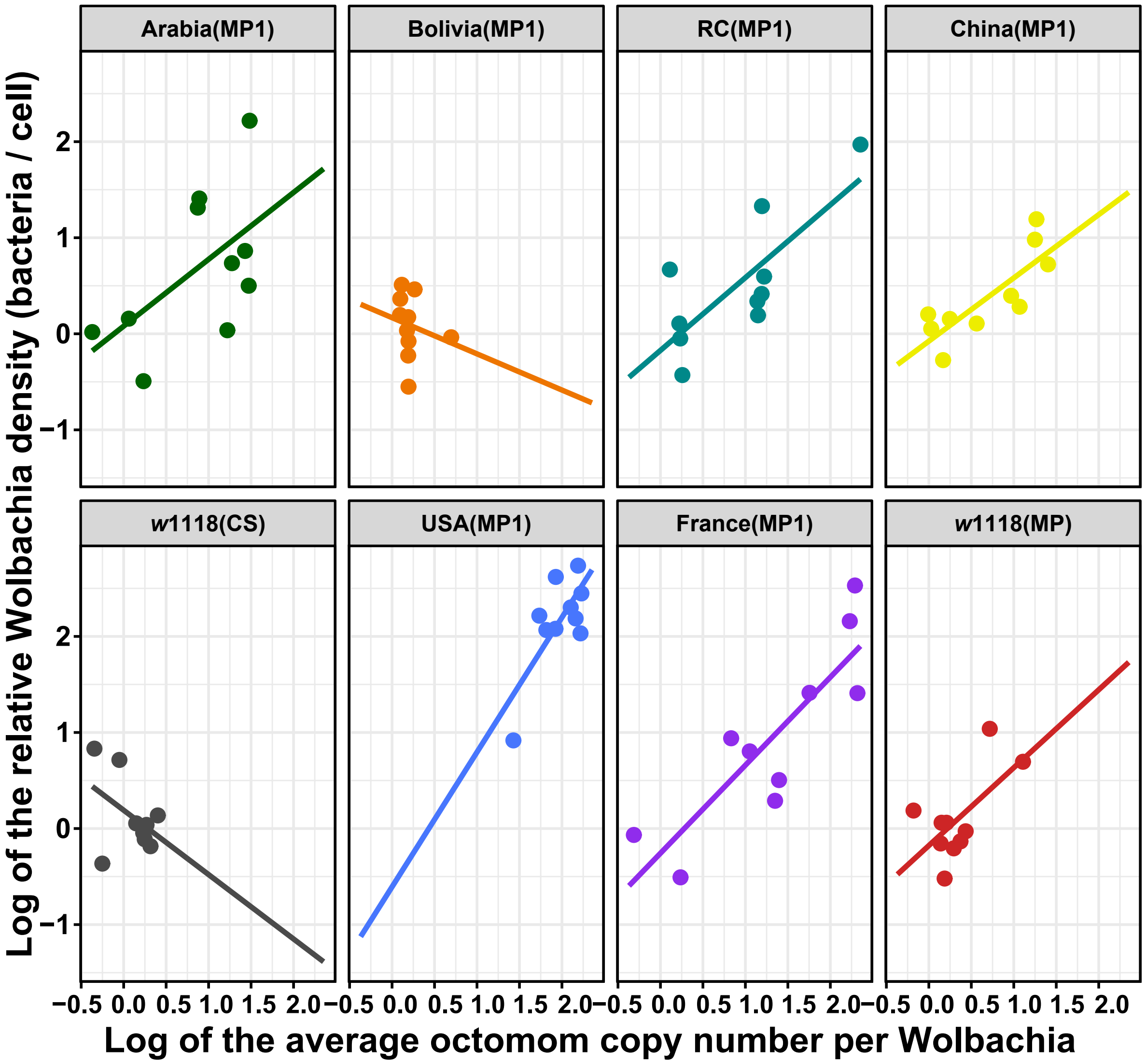


Arabia(MP1): 0.696x + 0.079; $R^2$ = 0.36; *P* = 0.085
Bolivia(MP1): -0.377x + 0.168; $R^2$ = 0.04; *P* = 0.5753
**RC(MP1): 0.758x -0.173; $R^2$ = 0.59; *P* = 0.009**
**China(MP1): 0.659x -0.077; $R^2$ = 0.66; *P* = 0.005**
W1118(wMelCS): -0.670x -0.191; $R^2$ = 0.19; *P* = 0.236
**USA(MP1): 1.4022x -0.607; $R^2$ = 0.54; *P* = 0.001**
**France(MP1): 0.916x – 0.257; $R^2$ = 0.73; *P* = 0.002**
**W1118(MP): 0.811x -0.177; $R^2$ = 0.40; *P* = 0.049**

**Figure s2:** Relationship between the relative density and the average octomom copy number per ***Wolbachia*** **in replicate lines at each timepoint of the new introgression experiment** (exp. #2).

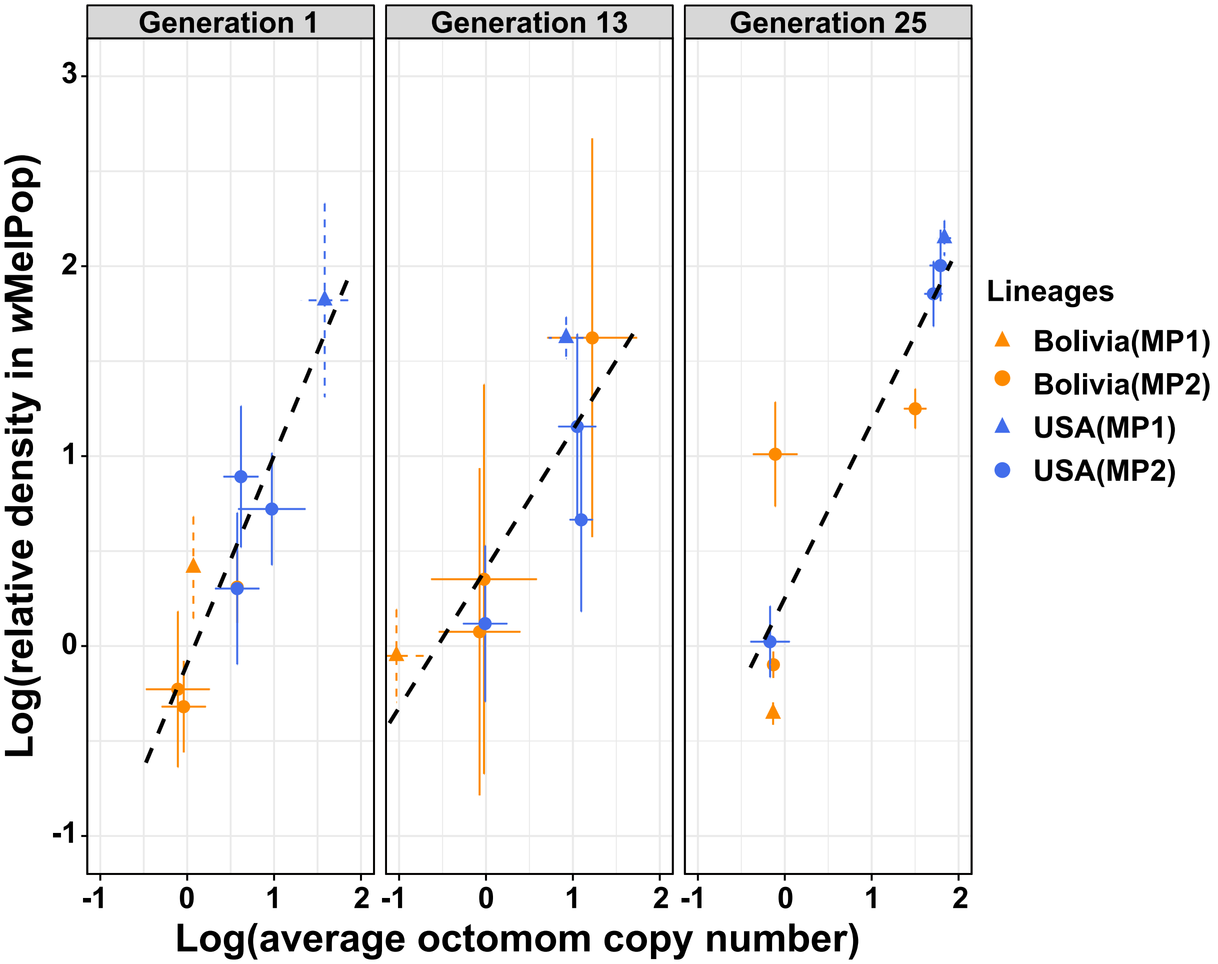


**Generation #1: 1.096x – 0.093; $R^2$ = 0.84; *P* = 0.001**
**Generation #13: 0.731x + 0.407; $R^2$ = 0.72; *P* = 0.008**
**Generation #25: 0.993x + 0.252; $R^2$ = 0.83; *P* = 0.002**

**<u>Figure s3:</u> Correlation between the relative density and the average octomom copy number per *Wolbachia* within each replicate line of the new introgression experiment** (exp. #2).

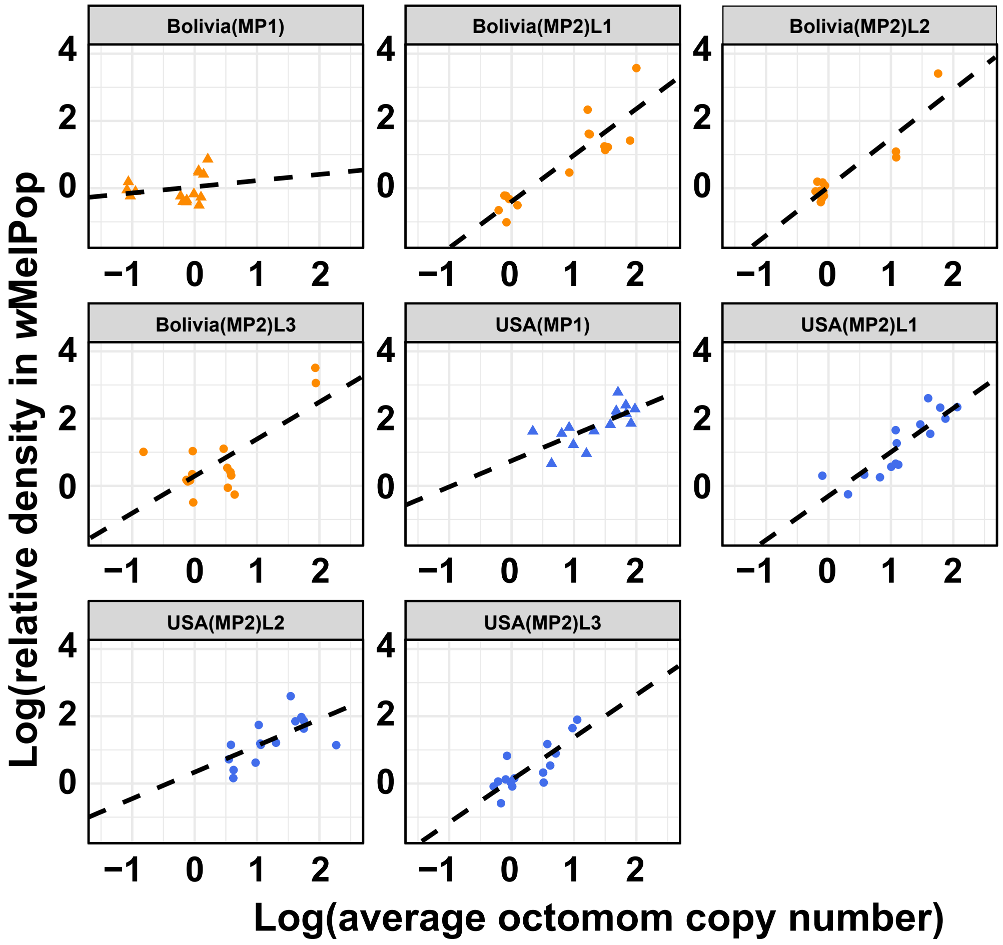


Bolivia(MP1): 0.176x + 0.030; $R^2$ = 0.044; $P$ = 0.455

**Bolivia(MP2)L1: 1.390x – 0.392; $R^2$ = 0.79; $P$ < 0.001**

**Bolivia(MP2)L2: 1.460x + 0.042; $R^2$ = 0.86; $P$ < 0.001**

**Bolivia(MP2)L1: 1.098x + 0.296; $R^2$ = 0.52; $P$ = 0.003**

**USA(MP1): 0.773x – 0.751; $R^2$ = 0.50; $P$ = 0.003**

**USA(MP2)L1: 1.312x – 0.317; $R^2$ = 0.75; $P$ < 0.001**

**USA(MP2)L2: 0.796x + 0.317; $R^2$ = 0.41; $P$ = 0.011**

**USA(MP2)L3: 1.260x + 0.112; $R^2$ = 0.66; $P$ < 0.001**

**<u>Figure s4:</u> Correlation between the relative density and the average octomom copy number per *Wolbachia* in replicate lines at each timepoint of the 'reciprocal crosses' experiment** (exp. #3).

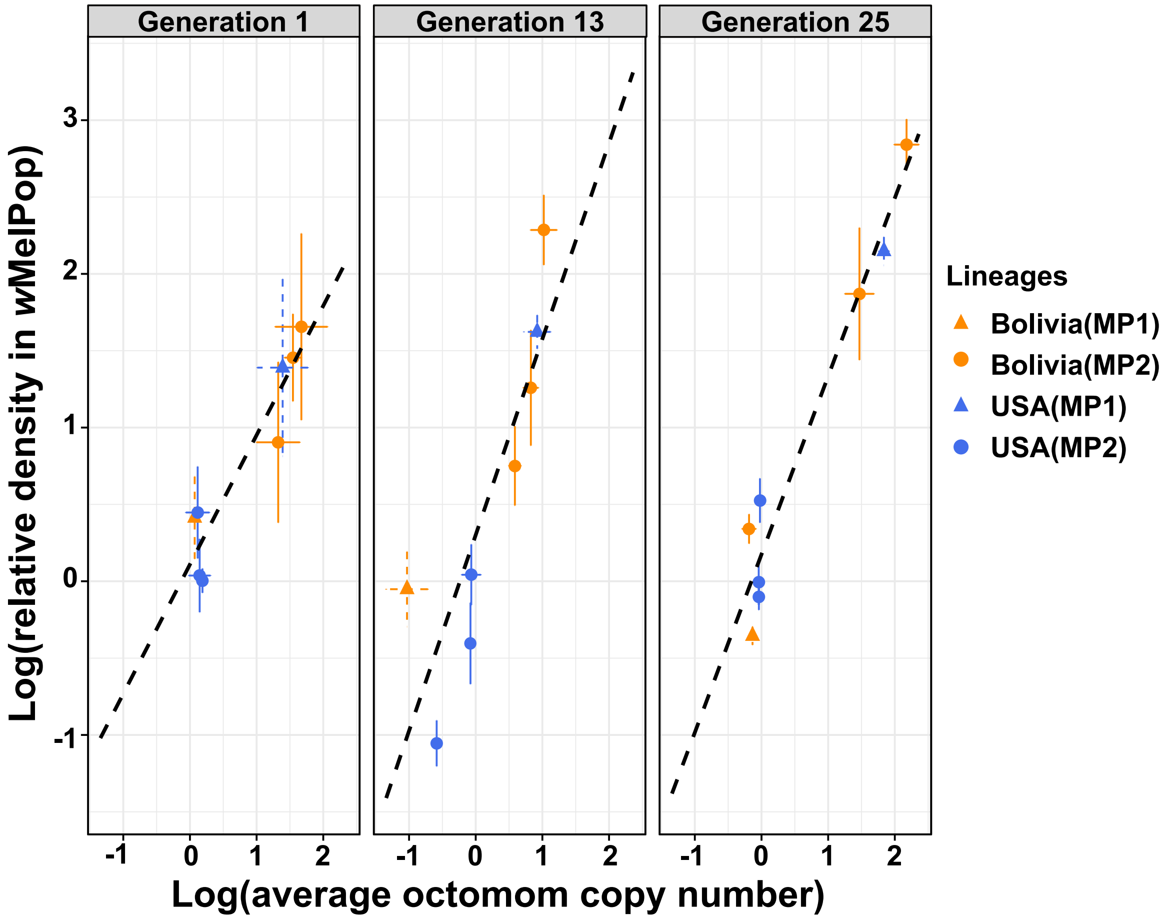


**Generation #1: 0.854x + 0.109; $R^2$ = 0.88 *P* < 0.001**
**Generation #13: 1.275x + 0.302; $R^2$ = 0.74; *P* = 0.006**
**Generation #25: 1.83x – 0.763; $R^2$ = 0.96; *P* < 0.001**

<u>Figure s5:</u> **Correlation between the relative density and the average octomom copy number per *Wolbachia* within each line of the 'reciprocal crosses' experiment** (exp. #3).

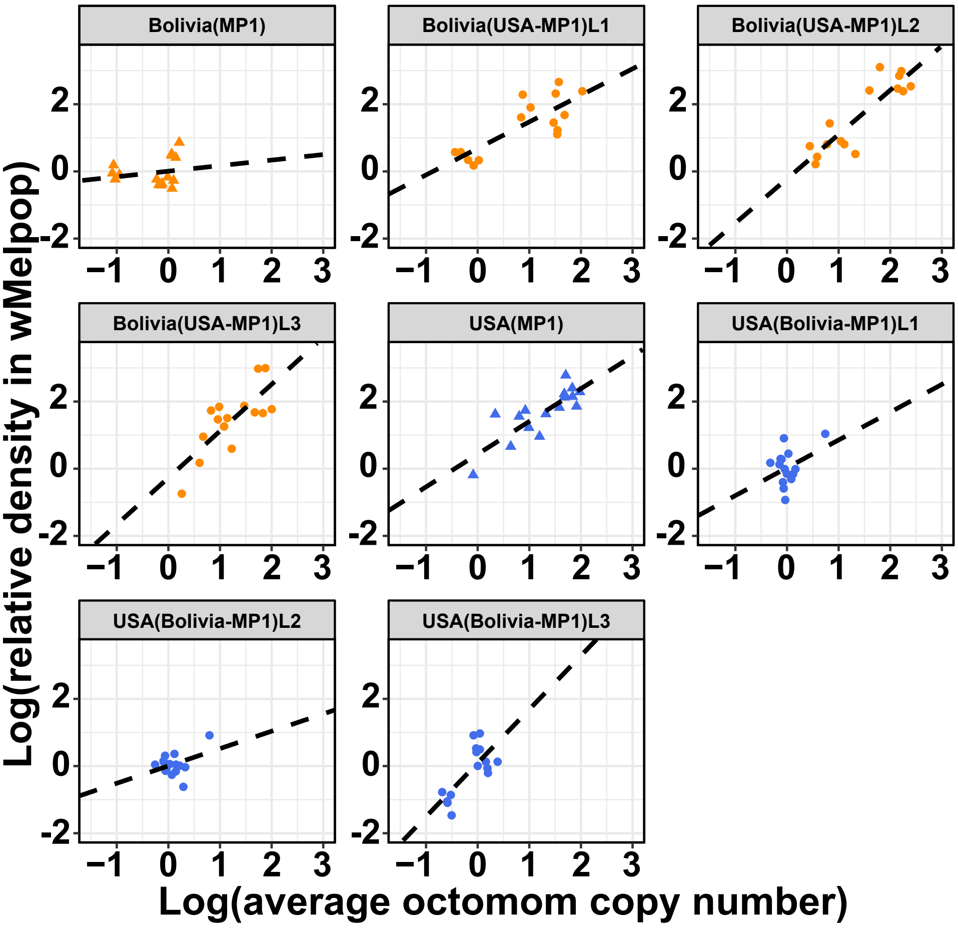


Bolivia(MP1): 0.176x + 0.030; $R^2$ = 0.04; *P* = 0.455
**Bolivia(USA-MP1)L1: 0.783 + 0.692; $R^2$ = 0.54; *P* < 0.001**
**Bolivia(USA-MP1)L2: 1.319x - 0.229; $R^2$ = 0.78; *P* < 0.001**
**Bolivia(USA-MP1)L3: 1.400x - 0.025; $R^2$ = 0.58; *P* < 0.001**
**USA(Bolivia-MP1): 0.773x – 0.751; $R^2$ = 0.50; *P* = 0.003**
USA(Bolivia-MP1)L1: 0.827x + 0.039; $R^2$ = 0.14; *P* = 0.173
USA(Bolivia-MP1)L2: 0.510x - 0.009; $R^2$ = 0.14; *P* = 0.167
**USA(Bolivia-MP1)L3: 1.612x + 0.084; $R^2$ = 0.54; *P* = 0.002**
